# Genetically encoded biosensors for branched-chain amino acid metabolism to monitor mitochondrial and cytosolic production of isobutanol and isopentanol in yeast

**DOI:** 10.1101/2020.03.08.982801

**Authors:** Yanfei Zhang, Sarah K. Hammer, Cesar Carrasco-Lopez, Sergio A. Garcia Echauri, José L. Avalos

## Abstract

Branched-chain amino acid (BCAA) metabolism can be harnessed to produce many valuable chemicals. Among these, isobutanol, which is derived from valine degradation, has received substantial attention due to its promise as an advanced biofuel. While *Saccharomyces cerevisiae* is the preferred organism for isobutanol production, the lack of isobutanol biosensors in this organism has limited the ability to screen strains at high throughput. Here, we use a transcriptional regulator of BCAA biosynthesis, Leu3p, to develop the first genetically encoded biosensor for isobutanol production in yeast. Small modifications allowed us to redeploy Leu3p in a second biosensor for isopentanol, another BCAA-derived product of interest. Each biosensor is highly specific to isobutanol or isopentanol, respectively, and was used to engineer metabolic enzymes to increase titers. The isobutanol biosensor was additionally employed to isolate high-producing strains, and guide the construction and enhancement of mitochondrial and cytosolic isobutanol biosynthetic pathways, including in combination with optogenetic actuators to enhance metabolic flux. These biosensors promise to accelerate the development of enzymes and strains for branched-chain higher alcohol production, and offer a blueprint to develop biosensors for other products derived from BCAA metabolism.

## Introduction

The metabolism of branched chain amino acids (BCAAs), including valine, leucine, and isoleucine, has been the subject of countless fundamental and applied studies^1–8^, and is central to the biosynthesis of many valuable products^9–16^. Among these products, branched-chain higher alcohols (BCHAs) including isobutanol, isopentanol, and 2-methyl-1-butanol are promising alternatives to the first-generation biofuel, ethanol. These alcohols exhibit superior fuel properties to ethanol, such as higher energy density, as well as lower vapor pressure and water solubility, which result in better compatibility with existing gasoline engines and distribution infrastructure^14^. The Co-Optimization of Fuels & Engines (Co-Optima) initiative sponsored by the United States Department of Energy (DOE) evaluated more than 400 compounds for their potential to enhance engine efficiency, and found isobutanol and blends of the three BCHAs to be in the top ten performers for boosted spark-ignition engines^17, 18^. Furthermore, BCHAs can be upgraded to jet fuels^19^, offering a source of renewable energy for air transportation. Co-Optima’s identification of these BCHAs as valuable renewable fuel compounds validates previous work to engineer microbes for the production of BCHAs, and encourages future work to this end.

The yeast *Saccharomyces cerevisiae* is a favored industrial host due to its resistance to phage contamination, ability to grow at low pH, and genetic tractability^20^. For BCHA production in particular, *S. cerevisiae* benefits from higher tolerance to alcohols than many bacteria^21^. In addition, BCHAs are naturally produced in *S. cerevisiae* as products of BCAA degradation. Accordingly, metabolic pathways for BCHA production have been engineered by rewiring BCAA biosynthesis and the Ehrlich degradation pathway (**Supplementary Figure 1**). The upstream BCAA biosynthetic pathway is comprised of three mitochondrially localized enzymes: acetolactate synthase (ALS, encoded by *ILV2*), ketol-acid reductoisomerase (KARI, encoded by *ILV5*), and dehydroxyacid dehydratase (DHAD, encoded by *ILV3*)^22^. Ilv2p, Ilv3p, and Ilv5p convert pyruvate to the valine precursor *α*-ketoisovalerate (*α*-KIV), which can be exported to the cytosol and converted to isobutanol via the downstream BCAA Ehrlich degradation pathway^23^, comprised of *α*-ketoacid decarboxylases (*α*-KDCs) and alcohol dehydrogenases (ADHs). Alternatively, *α*-KIV can be converted to α-ketoisocaproate (*α*-KIC) by sequential reactions carried out by *α*-isopropylmalate synthase (encoded by *LEU4* and *LEU9*), isopropylmalate isomerase (encoded by *LEU1*), and 3-isopropylmalate dehydrogenase (encoded by *LEU2*). The *LEU4* gene produces two forms of α-isopropylmalate synthase – a short form located in the cytosol, and a long form localized in mitochondria along with Leu9p^22^. The same Ehrlich degradation pathway converts *α*-KIC to isopentanol in the cytosol^23^. The fragmentation of BCHA biosynthesis between the mitochondria and cytosol presents an intracellular transport bottleneck, which has been addressed by compartmentalizing the downstream enzymes in mitochondria^24–28^ or re-locating the upstream enzymes in the cytosol^29–34^.

The rate at which researchers can introduce diversity into engineered strains greatly outpaces the rate at which strains can be tested for chemical production phenotypes. This challenge can be alleviated by the development of genetically encoded biosensors, which with sufficient sensitivity, specificity, and dynamic range, can enable high throughput screens or selections to identify variants with enhanced chemical productivity^35–40^. While a general alcohol biosensor that responds to isobutanol has been reported in *E.coli*^41, 42^, no biosensors for BCHAs have been reported in *S. cerevisiae*, which has prevented the development of high-throughput analysis of the production of this important class of biofuels in the preferred industrial host.

Many biosensors are based on transcription factors (TF) that activate expression of a reporter gene (such as a fluorescent protein) in response to elevated concentrations of a specific molecule, which may be the product of interest or a precursor^8, 37, 38, 43, 44^. Measuring levels of an intermediate intracellular metabolite, as opposed to a secreted product^42^, is advantageous because it allows the biosensor to capture the desired chemical production occurring in each individual cell rather than the extracellular product concentration, which reflects the average production of all cells in the fermentation. In *S. cerevisiae*, Leu3p, a dual-function TF, regulates several genes involved in BCAA biosynthesis^45^ (**Supplementary Figure 1**). Leu3p responds selectively to *α*-isopropylmalate (*α*-IPM), acting as a transcriptional activator in the presence of α-IPM and a transcriptional repressor in its absence. Because α-IPM is a byproduct of valine biosynthesis and an intermediate in leucine biosynthesis, its levels are correlated to the metabolic fluxes through both of these pathways and thus isobutanol and isopentanol production. This makes Leu3p an attractive TF with which to engineering a biosensor for BCHA biosynthesis in *S. cerevisiae*.

Here, we report the development and application of the first genetically encoded biosensors for BCAA metabolism and BCHA biosynthesis in *S. cerevisiae*. The biosensors are based on the α-IPM-dependent transcription factor function of Leu3p, which we use to control the expression of green fluorescent protein (GFP) as a reporter. Small differences in design permit the development of two distinct biosensors – one specific to isobutanol production and one specific to isopentanol production, which enable the development of high-throughput screens for both biosynthetic pathways. We apply these biosensors to isolate high-producing strains, identify mutant enzymes with enhanced activity, and construct biosynthetic pathways in both mitochondria and cytosol for the production of isobutanol and isopentanol. These biosensors have the potential to accelerate the development of strains to produce BCHAs as well as other products derived from BCAA metabolism.

## Results

### Design and construction of biosensors for branched-chain higher alcohol biosynthesis

The role of Leu3p as transcriptional regulator of branched-chain amino acid (BCAA) biosynthesis can be used to design genetically encoded biosensors for BCAA or branched-chain higher alcohol (BCHA) production in yeast. The dual transcriptional regulatory activity of Leu3p is mediated through its interaction with α-isopropylmalate (α-IPM), a key intermediate of BCAA biosynthesis^45^ (**Supplementary Fig. 1**). In its basal (*apo*) state, Leu3p acts as a transcriptional repressor of genes involved in BCAA biosynthesis (**Supplementary Fig. 1**). However, in the presence of α-IPM, Leu3p becomes a transcriptional activator of the genes it represses in its basal state. The α-IPM synthase Leu4p, and its paralog Leu9p, produce α-IPM from α-ketoisovalerate (α-KIV) in what amounts to the first committed step to leucine and isopentanol biosynthesis, and the branch point away from valine and isobutanol biosynthesis. This makes α-IPM both a byproduct of isobutanol (and valine), and a precursor of isopentanol (and leucine) (**Supplementary Fig. 1**). Thus, we reasoned that transcriptional activity from α-IPM-activated Leu3p could be used to monitor the metabolic flux toward either isobutanol or isopentanol production.

To construct a biosensor for isobutanol production, we introduced a copy of yeast-enhanced green fluorescent protein (yEGFP) under the control of the *LEU1* promoter (PLEU1), which is regulated by Leu3p, and a copy of a leucine-insensitive Leu4p. In yeast, leucine biosynthesis is negatively regulated by feedback inhibition of Leu4p by leucine (**Supplementary Fig. 1**). Thus, in the presence of leucine, α-IPM production decreases and the activity of Leu3p shifts from a transcriptional activator to a transcriptional repressor. Therefore, to obtain an isobutanol biosensor that functions independently of the intracellular levels of leucine, it is necessary to use a leucine-insensitive mutant of *LEU4*. To achieve this, we expressed a Leu4p mutant with its leucine regulation domain truncated (Leu4^1–410^) under the control of the constitutive *TPI1* promoter (PTPI1). Our results suggest that this truncated Leu4p dimerizes with the endogenous Leu4p, Leu4^WT^, forming a catalytically active Leu4^WT^/Leu4^1–410^ complex insensitive to feedback inhibition from leucine. The strain housing the isobutanol biosensor has a deletion in *LEU2*, causing α-IPM to accumulate as a byproduct depending on the metabolic flux to isobutanol biosynthesis, which proportionally enhances Leu3p-mediated expression of yEGFP. To reduce the background signal and prevent α-IPM accumulation from rapidly saturating the biosensor, we fused a proline-glutamate-serine-threonine-rich (PEST) protein degradation tag^46^ to the C-terminus of yEGFP (**Fig. 1a**). Thus, our biosensor for isobutanol biosynthesis consists of a simple construct containing PLEU1-yEGFP-PEST-TADH1 and PTPI1-*LEU4*^1-410^-TPGK1, in a *leu2Δ* strain.

**Figure. 1.**
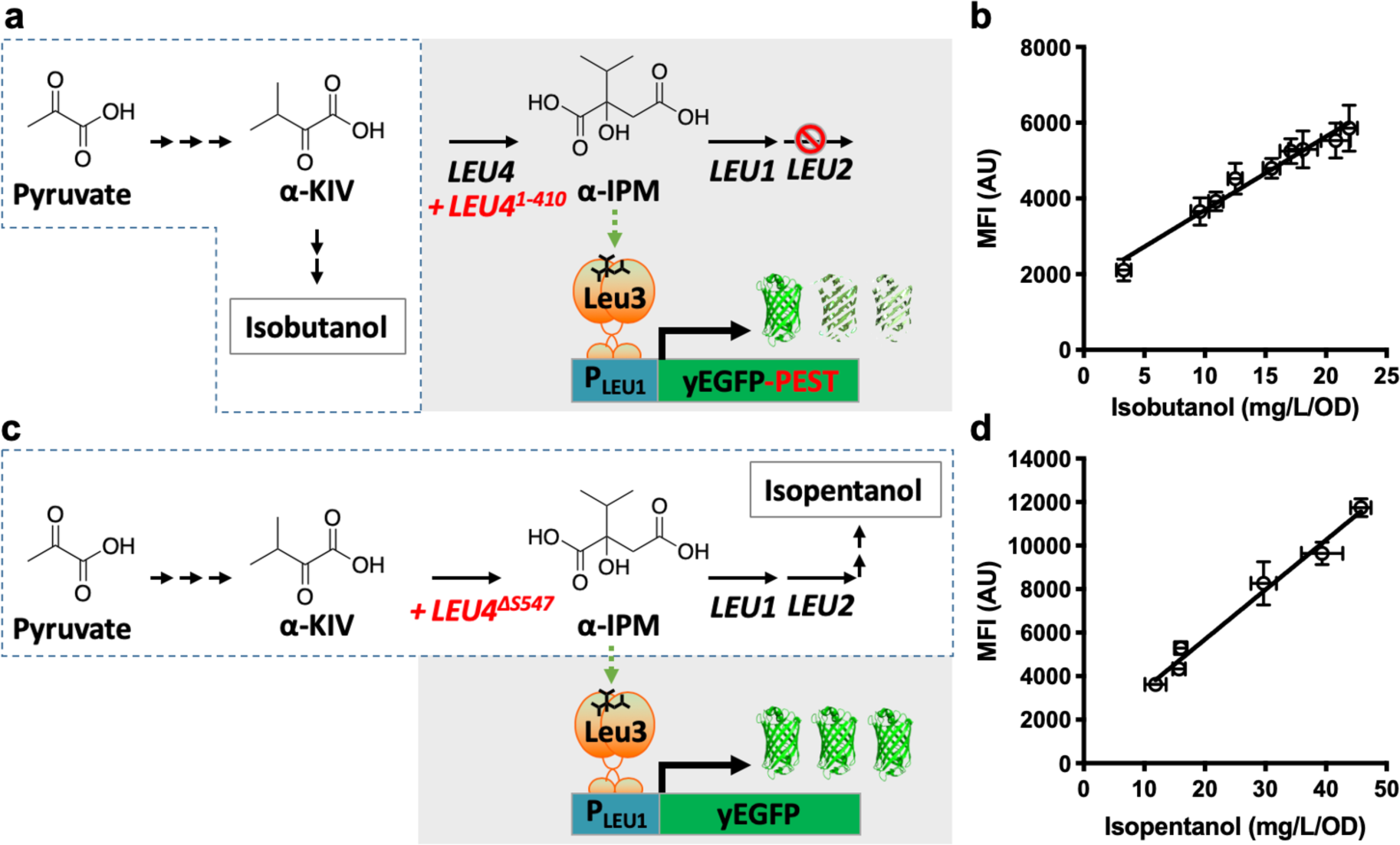
Design and characterization of Leu3-based biosensors for isobutanol and isopentanol. (a) Schematics of the isobutanol biosensor (gray box) and simplified isobutanol pathway (enclosed by dashed lines; each arrow represents one enzymatic step). The α-isopropylmalate (α-IPM), which activates Leu3p (green dashed arrow), is a byproduct of isobutanol biosynthesis, and accumulates due to deletion of *LEU2*. Thus, adding a PEST tag to destabilize yEGFP prevents early saturation of the biosensor signal. α-KIV, *α*-ketoisovalerate. (**b**) Correlation between specific isobutanol titers (mg/L/OD) produced by engineered strains and GFP median fluorescence intensity (MFI) from the isobutanol biosensor. Strains used (from left to right): YZy121, YZy230, YZy81, YZy231, YZy232, YZy233, YZy234, YZy235, and YZy236 (**Supplementary Table 1)**. (**c**) Schematics of the isopentanol biosensor (gray box) and simplified isopentanol pathway (enclosed by dashed lines; each arrow represents one enzymatic step). In this case, α-IPM is a precursor of isopentanol biosynthesis and does not accumulate, so the PEST tag on yEGFP is not required. (**d**) Correlation between specific isopentanol titers (mg/L/OD) produced by engineered strains and GFP median fluorescence intensity (MFI) from the isopentanol biosensor. Strains used (from left to right): SHy187, SHy188, SHy192, SHy159, SHy176, and SHy158 (**Supplementary Table 1**). The MFI for each strain was measured after 13 h of cultivation, during the exponential phase, and plotted against the corresponding specific isobutanol or isopentanol titers obtained after 48-h high-cell-density fermentations. Error bars represent the standard deviation of at least three biological replicates.

To characterize this isobutanol biosensor, we measured its response to key metabolites supplemented in the growth medium as well as to enhanced isobutanol biosynthesis in engineered strains. We found that the fluorescence emitted by the biosensor responds linearly to increasing concentrations of α-IPM (R^2^ = 0.96, **Supplementary Fig. 2a**) or α-KIV (R^2^ = 0.97, **Supplementary Fig. 2b**) supplemented in the media. We next examined the correlation between GFP fluorescence and isobutanol production in strains containing the biosensor and different constructions of the isobutanol pathway, resulting in varying levels of isobutanol production. We found that there is a strong linear correlation (R^2^ = 0.97) between GFP fluorescence and isobutanol biosynthesis (**Fig. 1b**), validating the applicability of this biosensor over at least an 11-fold range of isobutanol production.

The isobutanol biosensor can be adapted to generate an isopentanol biosensor by making only three modifications to the biosensor design and strain background. First, to produce isopentanol, it is necessary to have a strain that expresses *LEU2* (**Supplementary Fig. 1**). Because α-IPM is an intermediate metabolite of isopentanol biosynthesis (**Supplementary Fig. 1**), intracellular concentrations of α-IPM are expected to be substantially lower in strains engineered to make isopentanol than in *leu2Δ* strains engineered to make isobutanol, in which α-IPM may accumulate as a byproduct. Thus, when sensing α-IPM in an isopentanol-producing strain, the need to improve the biosensor limit of detection is greater than the need to prevent its GFP signal from saturating. To achieve this, as a second modification to the isobutanol biosensor, we removed the PEST-tag from the yEGFP reporter. Lastly, we found that deleting the endogenous genes encoding for α-IPM synthases (*LEU4* and *LEU9*) reduces the background noise of the biosensor. However, this requires replacing Leu4^1-410^, which depends on endogenous *LEU4* for activity, with another leucine-insensitive Leu4p mutant. We chose to employ a Leu4p harboring a Ser547 deletion (Leu4^ΔS547^)^47^, which is active even in the absence of wild type *LEU4*^48^ (**Fig. 1c**). To validate the efficacy of this design, we measured the fluorescence of several strains containing this biosensor and different constructions of the isopentanol biosynthetic pathway, resulting in varying levels of isopentanol production. We found a strong linear correlation (R^2^ = 0.98) between the fluorescence emitted by the biosensor and isopentanol production, spanning at least a 4-fold range of isopentanol production (**Fig. 1d**). Therefore, a construct containing PLEU1-yEGFP-TADH1 and PTPI1-*LEU4^ΔS547^*-TPGK1 in a *leu4*, *leu9, LEU2* strain results in a functional biosensor for isopentanol production.

### Using the isobutanol biosensor to isolate high isobutanol-producing strains

We next applied the isobutanol biosensor to isolate the highest isobutanol-producers from a library of strains containing random combinations of genes from the mitochondrial isobutanol biosynthesis pathway^24^. Equal molar amounts of cassettes containing either the upstream *ILV* genes (*ILV2, ILV3,* and *ILV5*), the downstream Ehrlich pathway (KDC and ADH), or the full isobutanol pathway, all designed to randomly integrate into YARCdelta5 *δ*-sites^24, 49^, were pooled and used to transform strain YZy81, containing the isobutanol biosensor in the *HIS3* locus (**Supplementary Tables 1 and 2**). The transformed population was subjected to two rounds of fluorescence activated cell sorting (FACS, see Methods) followed by fermentations with 24 randomly picked colonies from each population (unsorted, first round sorted, and second round sorted). For each of the three populations analyzed, we calculated the average isobutanol titer **(Fig. 2a**) and assigned each colony to a titer interval in accordance with its isobutanol production **(Fig. 2b**). The average titers of the analyzed colonies increase after each round of sorting. Most colonies (16 out of 24) from the unsorted population produce between 100 mg/L and 300 mg/L isobutanol, which is similar to the titer of the host strain YZy81 (179 ± 30 mg/L), while only one out of the 24 screened colonies from the unsorted population produces more than 500 mg/L isobutanol. The majority of colonies selected after one round of sorting produce between 300 mg/L and 600 mg/L. After the second round of sorting, 11 of the 24 randomly chosen colonies produce more than 600 mg/L isobutanol, well above the titers achieved by any of the colonies from the unsorted population (**Fig. 2b**). These results demonstrate that our biosensor enables high-throughput screening with FACS to isolate high-isobutanol producers from mixed strain populations.

**Figure 2.**
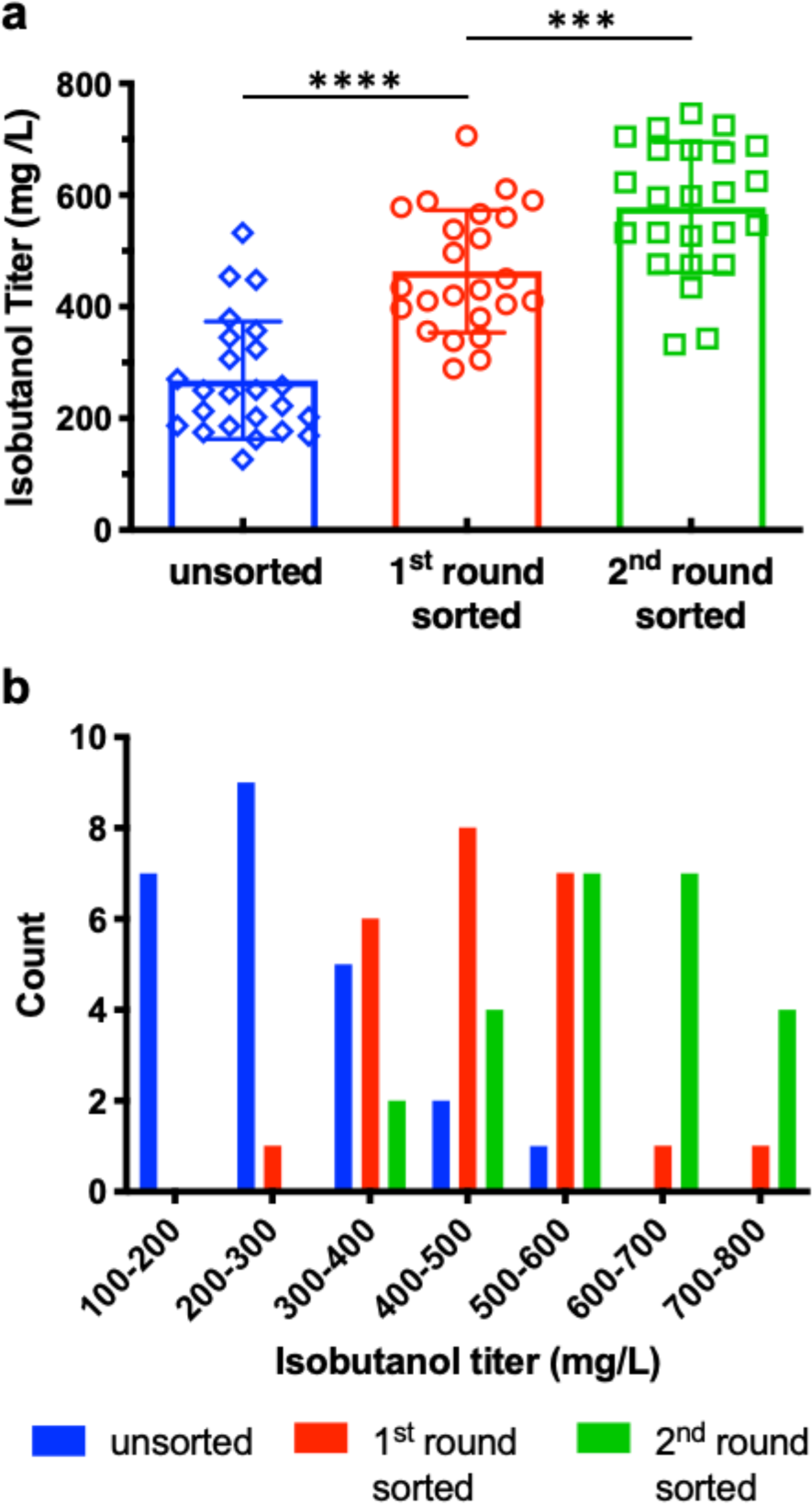
Applying the isobutanol biosensor for high-throughput screening of strains with the mitochondrial isobutanol biosynthetic pathway. (**a**) Isobutanol titers of 24 colonies randomly selected from unsorted (blue), first round sorted (red), or second round sorted (green) populations derived from a library of strains transformed with constructs overexpressing different combinations of the mitochondrial isobutanol pathway. Error bars represent the standard deviation of the titers of the 24 colonies analyzed. A two-tailed *t*-test was used to determine the statistical significance of the differences in isobutanol titers of analyzed strains from each population; *** *P* ≤ 0.001, **** *P* ≤ 0.0001. (**b**) Histogram showing the number of colonies exhibiting different ranges of isobutanol production, out of the same 24 randomly selected strains from the unsorted (blue), first round sorted (red), or second round sorted (green) populations shown in (a).

### Applying the isobutanol biosensor to identify ILV6 mutants insensitive to valine inhibition

We next applied our isobutanol biosensor to identify mutants of the acetolactate synthase regulatory protein, encoded by *ILV6,* with reduced feedback inhibition from valine. Ilv6p localizes to mitochondria and interacts with acetolactate synthase, Ilv2p, both enhancing its activity and bestowing upon it feedback inhibition by valine^50, 51^ (**Fig. 3a**). Mutations identified in the Ilv6p homolog from *Streptomyces cinnamonensis* (IlvN) that confer resistance to valine analogues^52^ informed the design of Ilv6p^V90D/L91F^, which has been shown to retain Ilv2p activation, have reduced sensitivity to valine inhibition, and improve isobutanol production^25, 26^. However, these mutations are likely not unique or optimal; if alternative mutations exist, some potentially conferring higher Ilv2p/Ilv6p complex activity, we hypothesized that we could find them with our biosensor.

**Figure 3.**
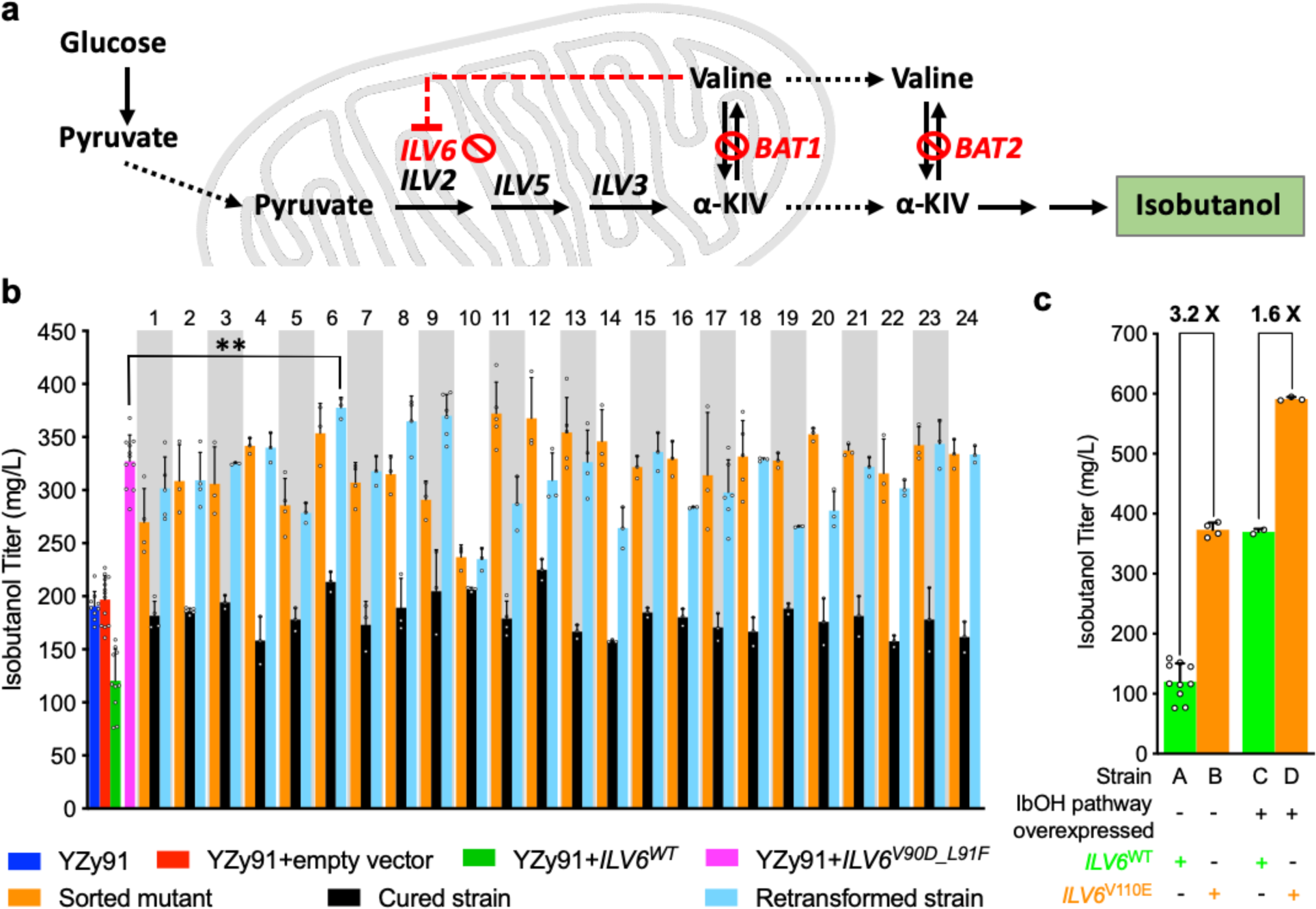
High-throughput screen for metabolically hyperactive Ilv6p mutants using the isobutanol biosensor. (**a**) Schematic representation of the genotype of the basal strain, YZy91, used for high-throughput screening of Ilv6p mutants with enhanced activity in the presence of valine; *α*-KIV: *α*-ketoisovalerate. (**b**) Isobutanol titers after 48h fermentations of sorted strains with unique *ILV6* mutant sequences (orange), plasmid-cured derivatives (black), and strains obtained from retransforming YZy91 with each unique plasmid isolated from the corresponding sorted strain (cyan). Titers obtained from YZy91 with (red) or without (blue) an empty vector, or transformed with plasmids containing *ILV6^WT^* (green) or *ILV6^V90D_L91F^* (magenta), are shown as controls. The numbers on the upper x-axis are used to identify each of the strains harboring screened mutants with unique *ILV6* sequences. *ILV6* mutants 1 – 10 were isolated from the sorted wild-type *ILV6* (*ILV6^WT^*) mutagenesis library. *ILV6* mutants 11 – 24 were isolated from the sorted *ILV6^V90D_L91F^* mutagenesis library. A two-tailed *t*-test was used to determine the statistical significance of the difference in isobutanol titers between YZy91 with *ILV6^V90D_L91F^* (magenta) and *ILV6^V110E^* (*ILV6* mutant #6). ** *P* ≤ 0.01. (**c**) Isobutanol titers of YZy91 transformed with CEN plasmids containing *ILV6^WT^* (Strain A) or *ILV6^V110E^* (*ILV6* mutant #6, Strain B) compared to titers from a strain overexpressing the mitochondrial isobutanol pathway (YZy363) transformed with CEN plasmids containing *ILV6^WT^* (Strain C) or *ILV6^V110E^* (*ILV6* mutant #6, Strain D). Error bars represent the standard deviation of at least three biological replicates.

To find Ilv6p mutants insensitive to valine inhibition, we used our isobutanol biosensor to isolate strains with the highest fluorescence in the presence of valine from randomly mutagenized *ILV6* libraries. We introduced the isobutanol biosensor into a strain containing *BAT1* and *BAT2* deletions to eliminate interconversion of valine and *α*-KIV, which would interfere with our biosensor signal, resulting in strain YZy91 (**Supplementary Table 1**). YZy91 also lacks the native *ILV6*, eliminating endogenous valine-inhibition of Ilv2p (**Fig. 3a**). Two libraries, generated via error-prone PCR of either the wild-type *ILV6* or the *ILV6^V90D/L91F^* variant, were used separately to transform YZy91. After culturing the two resulting yeast libraries in media containing excess valine (See Methods) and three rounds of FACS, we measured the isobutanol production of 24 colonies randomly picked from each sorted library. All 48 colonies randomly selected after the third round of sorting show higher biosensor signals in the presence of valine than the strain overexpressing wild-type *ILV6* (**Supplementary Fig. 3a**). More importantly, all 48 colonies screened produce higher isobutanol titers than the strain overexpressing the wild-type *ILV6* (**Supplementary Fig. 3b**), demonstrating that our biosensor can identify strains with improved isobutanol production without false positives.

To determine whether the improvements in isobutanol titers are caused by mutation(s) to *ILV6*, we genotyped and phenotyped the hyper-productive strains. We first cured each strain of its plasmid containing an *ILV6* variant by 5-fluoroorotic acid (5-FOA) counterselection (see Methods), while simultaneously isolating and sequencing the plasmids from each of the 48 strains. We found that 10 (out of 24) of the plasmids from the sorted wild-type *ILV6* mutagenesis library, and 14 (out of 24) of the plasmids from the sorted *ILV6^V90D/L91F^* mutagenesis library have unique *ILV6* sequences (**Supplementary Table 3**). Finally, we retransformed the parental strain (YZy91) with each of the 24 unique plasmids, and carried out fermentations with the sorted, cured, and retransformed strains (**Fig. 3b**). All of the cured strains produce isobutanol at titers similar to that of the parent strain, and most of the retransformed strains exhibit isobutanol titers comparable to, or higher than, those of the originally sorted stains. These phenotypes are reflected in the fluorescence intensity measurements for each strain, demonstrating the reliability of the biosensor (**Supplementary Fig. 4**). The few phenotypic differences observed between the originally sorted and retransformed strains are likely caused by unknown background mutations in the sorted strains. Nonetheless, our results show that all of the *ILV6* mutants isolated with our biosensor improve isobutanol production.

Our biosensor allowed us to identify a number of new *ILV6* mutations that enable at least the same level of isobutanol production as the previously identified valine-insensitive mutant, *ILV6^V90D/L91F^* (**Supplementary Table 3**). Among the 24 retransformed strains tested, the strain harboring *ILV6* mutant *#6*, which has a single mutation at Val110 (V110E), produces the most isobutanol (378 mg/L ± 10 mg/L), representing a 3.2-fold increase over the strain overexpressing wild-type *ILV6* (**Fig. 3b**). This strain also shows a small but statistically significant *(P* = 0.0055) improvement in isobutanol production compared to the strain harboring the previously identified *ILV6^V90D/L91F^* mutant. When we introduce this *ILV6^V110E^* variant into a strain overexpressing the mitochondrial isobutanol biosynthetic pathway, the production is 1.6 times higher than with the equivalent strain harboring wild-type *ILV6* (**Fig. 3c**). Mapping the mutations found in our isolated variants onto the crystal structure of the bacterial Ilv6p homologue IlvH revealed that most of the mutations are located in the regulatory valine binding domain (**Supplementary Fig.5**). Interestingly, Val110 is located inside the putative valine binding site, interacting with the L91 residue mutated in *ILV6^V90D/L91F^* (**Supplementary Fig.5b**). Several other mutations that increase Ilv2p activity are also located inside the valine binding pocket (N86, V90, L91, and N104), consistent with other mutations previously reported to make Ilv2p or its homologues insensitive to valine inhibition^25, 26, 52–54^. Our isobutanol biosensor is able to identify new *ILV6* mutations derived from mutagenizing the wild-type *ILV6* that improve isobutanol production (mutants #1 – #10 in **Supplementary Table 1**); however, none of the strains sorted from the *ILV6^V90D/L91F^* mutagenesis library (mutants #11 – #24 in **Supplementary Table 1**) show a significant improvement in isobutanol production relative to the strain overexpressing *ILV6^V90D/L91F^*. This could be because *ILV6^V90D/L91F^* enhances Ilv2p activity to an extent that is near the maximum achievable through Ilv6p engineering alone, or because the pathway bottleneck is no longer at the Ilv2p metabolic step after *ILV6^V90D/L91F^* is overexpressed.

### Applying the isopentanol biosensor to identify highly active LEU4 mutants

We next applied our isopentanol biosensor to identify mutant *LEU4* enzymes with enhanced activity. Our isopentanol biosensor is designed with a *LEU4* mutant previously shown to have reduced sensitivity to leucine inhibition^47, 55^. However, engineering a Leu4p variant that is not only insensitive to leucine inhibition, but also as catalytically active as possible is important when engineering strains for isopentanol production. We constructed two random mutagenesis libraries of *LEU4* with error-prone PCR: one in which the entire *LEU4* open reading frame (ORF) is mutagenized, and another in which only the regulatory domain (residues 430-619) is mutagenized. These libraries were used separately to transform YZy148, a strain containing a modified isopentanol biosensor lacking *LEU4^ΔS5^*^47^, and lacking endogenous *BAT1*, *LEU4*, and *LEU9* genes (**Supplementary Tables 1 and 2**). Analyzing cell cultures grown in the presence of leucine with flow cytometry revealed two major sub-populations in the library derived from mutagenizing the full-length *LEU4* (**Fig. 4a**). The larger population overlaps with the signal from the empty vector control, suggesting that most mutations are deleterious to Leu4p activity, and thus *α*-IPM synthesis. However, a smaller population retains activity, with a substantial fraction exhibiting higher GFP fluorescence intensity than the wild-type *LEU4*. In contrast, the library derived from mutagenizing only the regulatory domain of *LEU4* is dominated by a single population with a higher GFP signal than wild-type *LEU4*, suggesting that many mutations to the regulatory domain enhance Leu4p enzymatic activity in the presence of leucine, and few cause complete loss of activity.

**Figure 4.**
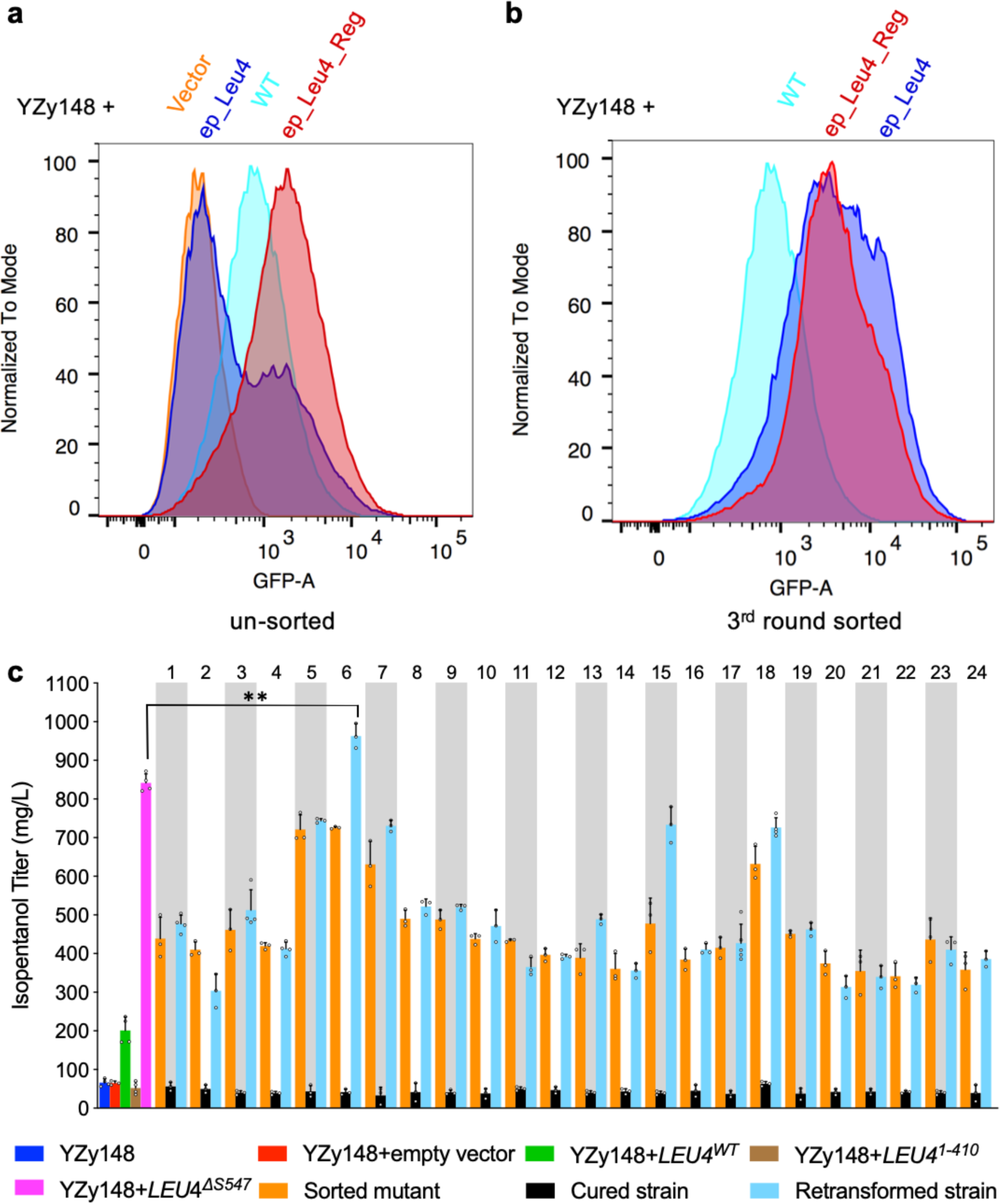
High-throughput screen for metabolically hyperactive Leu4p variants using the isopentanol biosensor. (**a**) Flow cytometry analysis of yeast libraries containing *LEU4* variants generated with error-prone PCR-based mutagenesis of the entire *LEU4* open reading frame (ep_Leu4, blue) or the *LEU4* regulatory domain (ep_Leu4_Reg, red), compared to control strains containing an empty vector (Vector, orange) or wild-type *LEU4* (WT, cyan). Cell populations were normalized to mode for comparison. (**b**) Flow cytometry analysis of the same libraries depicted in (a) after three rounds of FACS. (**c**) Isopentanol titers after 48h fermentations of sorted strains with unique *LEU4* sequences (orange), plasmid-cured derivatives (black), and strains obtained from retransforming YZy148 with each unique plasmid isolated from the corresponding sorted strain (cyan). Titers obtained from the basal strain YZy148 (*leu4Δ leu9Δ bat1Δ LEU2*) with (red) or without (blue) an empty vector, or transformed with plasmids containing wild-type *LEU4* (*LEU4^WT^*, green), *LEU4^1-410^* (brown), or *LEU4^ΔS547^* (magenta), are shown as controls. The numbers on the upper x-axis identify each strain harboring a unique *LEU4* sequence. *LEU4* mutants 1 – 13 are derived from mutagenizing the entire *LEU4* ORF; and mutants 14 – 24 from mutagenizing the *LEU4* regulatory domain. Error bars represent the standard deviation of at least three biological replicates. A two-tailed *t*-test was used to determine the statistical significance of the difference between isopentanol titers achieved by YZy148 transformed with a plasmid containing *LEU4^ΔS547^* (magenta), or *LEU4* mutant #6 (LEU4E86D_K191N_K374R_A445T_S481R_N515I_A568V_S601A). ** P ≤ 0.01.

We proceeded to sort the cells with the brightest GFP signal in each library. After three rounds of FACS, both sorted populations display increased median fluorescence intensity (**Fig. 4b**). Among 48 randomly selected colonies (24 from each sorted library, **Supplementary Fig. 6**), we found 24 containing unique *LEU4* variants (13 of which are derived from mutagenizing the entire *LEU4* gene, **Supplementary Table 4**). The sorted strains containing these variants produce between 375 mg/L and 725 mg/L isopentanol, which represent 1.8- to 3.6-fold improvements over the stain overexpressing the wild-type *LEU4* (**Fig. 4c)**. Taking the same approach as for the *ILV6* variants above, we cured each strain from its *LEU4*-containing plasmid and retransformed the parent strain (YZy148) with the same plasmids to compare fluorescence intensity (**Supplementary Fig. 7**) and isopentanol production (**Fig. 4c**) between the sorted, cured, and retransformed strains. All of the cured strains display GFP fluorescence and isopentanol titers similar to those of the parent strain (YZy148). Moreover, all of the retransformed strains (except strain with mutant #2) exhibit isopentanol titers and GFP fluorescence similar to, or higher than (strains with mutants #6 and #15), their originally sorted counterparts. The strain retransformed with mutant #6 achieves the highest isopentanol titer (963 mg/L ± 32 mg/L), which is 4.8-times higher than the titer achieved by overexpressing the wild-type *LEU4*, and significantly higher than the titer obtained with the *LEU4^ΔS547^* mutant previously described^48^ (842 mg/L ± 23 mg/L; *P*-value = 0.0021).

Our isopentanol biosensor made it possible to identify several new mutations in *LEU4* that may contribute to reduced regulation, enhanced catalytic activity, or both. Five variants (*LEU4* mutants #5, #14, #18, #20, and #24) have single mutations (**Supplementary Table 4**), indicating that the corresponding changes in residues (H541R, Y485N, Y538N, V584E, and T590I, respectively) are responsible for the observed enhancement. All these mutations are located in the regulatory domain of Leu4p (**Supplementary Fig. 8**). Mutating residue H541 has been previously reported to make Leu4p resistant to Zn^2+^-mediated inactivation by CoA^47^. The other four mutations are located inside (Y538N, V584E, and T590I) or in the vicinity (Y485N) of the leucine binding site, suggesting that they result in decreased sensitivity to leucine inhibition. Although we have not yet elucidated the possible impact that individual mutations found in variants containing two or more mutations have on leucine sensitivity or catalysis, seven residues (K51, Q439, F497, N515, V584, D578, and T590) are substituted in more than one variant and are mutagenized in 14 of the 19 variants containing two or more mutations (**Supplementary Tables 4 and 5**), suggesting they are involved in enhancing Leu4p activity. All of these sites, except K51, are also located in the regulatory domain (**Supplementary Fig. 8**), making it likely that they are also involved in reducing regulatory inhibition of Leu4p. Most of them are located inside or in the vicinity of the regulatory leucine binding site. Remarkably, N515, located inside the leucine binding site, is substituted in five of the 24 variants we identified, including *LEU4* mutant #6, which is the variant that produces most isopentanol. On the other hand, Q439 is far from the leucine binding site but close to H541, suggesting it might be involved in Zn^2+^-mediated CoA inactivation of Leu4p. Careful biochemical characterization of each mutation identified in our screens will be required to elucidate their roles, if any, in Leu4p activity enhancement. Nevertheless, our biosensor was instrumental in identifying 12 previously unreported residues that enhance isopentanol production when mutated, and which are likely involved in Leu4p regulation by leucine or CoA, and/or Leu4p catalytic activity.

### Biosensor-assisted cytosolic isobutanol pathway engineering

While all previous isobutanol-producing strains in this study are engineered with the isobutanol pathway compartmentalized in mitochondria^24^, cytosolic pathways have also been developed^32–34, 56^. Interested in demonstrating that our isobutanol biosensor is useful for enhancing both mitochondrial and cytosolic isobutanol pathways, we employed our biosensor to construct and optimize an isobutanol biosynthetic pathway entirely in the yeast cytosol, comprised of heterologous ALS, KARI, and DHAD (*ILV*) genes. We deleted *ILV3* from strain YZy121 containing the isobutanol biosensor (**Supplementary Table 1**) to eliminate mitochondrial contribution to *α*-KIV synthesis and ensure that the biosensor signal and isobutanol production are derived only from cytosolic activity (**Fig. 5a**). We additionally deleted *TMA29*, encoding an enzyme with acetolactate reductase activity that competes with KARI, producing the by-product 2,3-dihydroxy-2-methyl butanoate (DH2MB)^57, 58^. We transformed the resulting strain (YZy443) with integration cassettes containing single copies of *Bs_alsS*, encoding an ALS from *Bacillus subtilis*^59^, *Ec*_*ilvC^P2D1-A1^*, encoding an engineered NADH-dependent KARI from *E. coli*^60^, and *Ll*_*ilvD*, encoding a DHAD from *Lactococcus lactis* (**Fig. 5a**). Because *Ll*_IlvD requires iron-sulfur (Fe/S) clusters as a cofactor, we also overexpressed *AFT1*, a transcription factor involved in iron utilization and homeostasis^61^. Together, these genes encode a metabolic pathway to convert pyruvate to *α*-KIV in the yeast cytosol, which can then be converted to isobutanol by endogenous KDCs and ADHs, naturally located in the cytosol (**Fig. 5a**).

**Figure 5.**
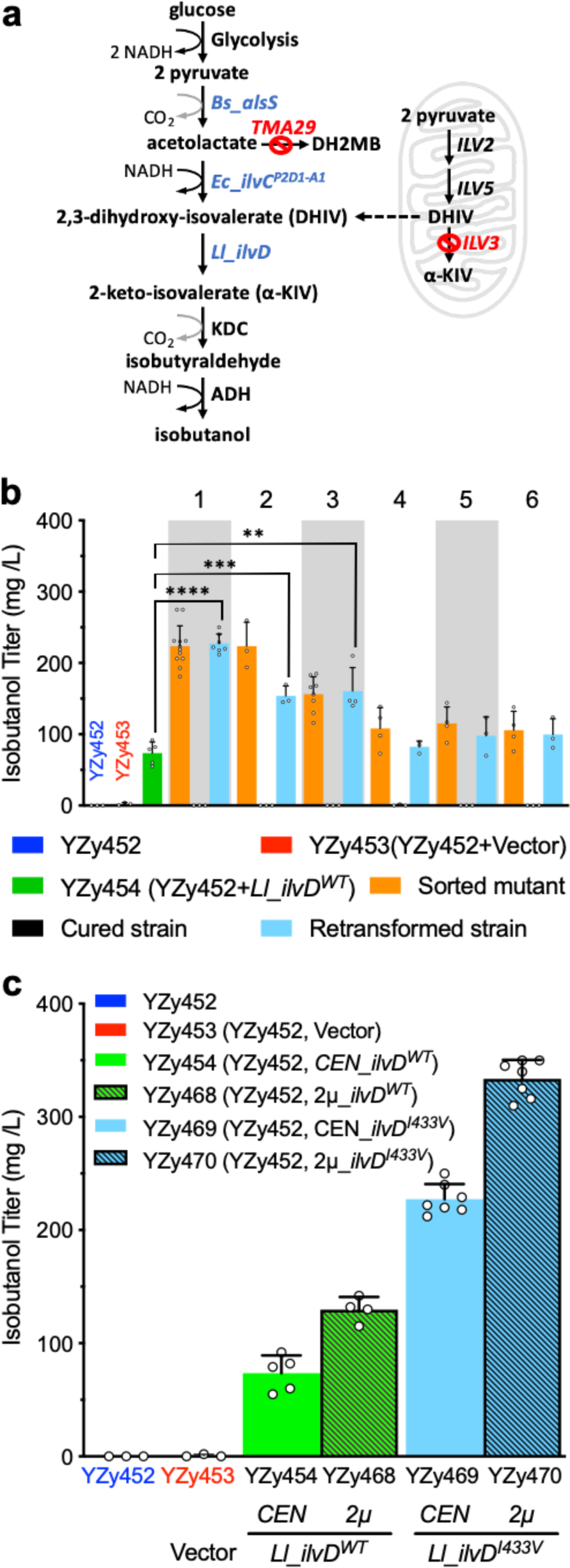
Biosensor-assisted cytosolic isobutanol pathway engineering. (**a**) Schematic of the engineered cytosolic isobutanol pathway. *Bs_alsS*: acetolactate synthase (ALS) from *Bacillus subtilis*; *Ec_ilvC^P2D1-A1^*: NADH-dependent ketol-acid reductoisomerase (KARI) variant from *E. coli*; *Ll_ilvD*: dihydroxyacid dehydratase (DHAD) from *Lactococcus lactis*; DH2MB: 2,3-dihydroxy-2-methyl butanoate. (**b**) Isobutanol titers after 48h fermentations in 15% glucose of sorted strains with unique *Ll_ilvD* sequences (orange), plasmid-cured derivatives (black), and strains obtained from retransforming YZy452 with each unique plasmid isolated from the corresponding sorted strain (cyan). The numbers on the upper x-axis identify each of the strains harboring screened mutants with unique *Ll_ilvD* sequences. Titers from YZy452 (containing the cytosolic isobutanol pathway with extra copies of *Ec_IlvC^P2D1-A1^*) with (red) or without (blue) an empty vector, or transformed with a plasmid containing wild type *Ll_ilvD* (green) are shown as controls. A two-tailed *t*-test was used to determine the statistical significance of the difference between isobutanol titers achieved by YZy452 transformed with a plasmid containing *Ll_ilvD^WT^* (green), or *Ll_ilvD* mutant #1 (*Ll_ilvD^I433V^*), or mutant #2 (*Ll_ilvD^V12A, S189P, H439R^*), or mutant #3 (*Ll_ilvD^K535R^*). ** *P* ≤ 0.01, *** *P* ≤ 0.001, **** *P* ≤ 0.0001. **(c)** Isobutanol titers of the basal stain YZy452 (blue) transformed with an empty vector (red), *Ll_ilvD^WT^* (green), or *Ll_ilvD^I433V^* (cyan), using CEN (solid) or 2μ (striped) plasmids. Isobutanol production was measured after 48h fermentations in 15% glucose. Error bars represent the standard deviation of at least three biological replicates.

We first applied our biosensor to find strains with increased levels of *Ec_ilvC^P2D1-A1^* expression to enhance isobutanol production in the cytosol. The baseline strain (YZy449) contains a single copy integration of the cytosolic isobutanol pathway, in which *Bs_alsS* and *Ec_ilvC^P2D1-A1^* are expressed under the control of strong constitutive promoters (PTEF1 and PTDH3, respectively), and *Ll_ilvD* is expressed by the galactose-inducible and glucose-repressible promoter PGAL10. Given that the catalytic rate constant (*k*cat) of *Bs_*AlsS is about 28 times higher than that of *Ec*_IlvC^P2D1-A1^ (**Supplementary Table 6**), we hypothesized that overexpressing additional copies of *Ec_ilvC^P2D1-A1^* would compensate for this difference and improve isobutanol production. Thus, we used our isobutanol biosensor to find the best producers from a library of strains containing additional copies of *Ec_ilvC^P2D1-A1^*. We randomly integrated a cassette containing two copies of *Ec_ilvC^P2D1-A1^* (pYZ206) to YARCdelta5 *δ*-sites of YZy449 (**Supplementary Tables 1 and 2**), and then used our biosensor to sort for transformants with the highest fluorescence intensity. After three rounds of FACS, we selected 20 random colonies and measured their isobutanol production. Both FACS and isobutanol fermentations where carried out in galactose-containing media to induce expression of *Ll_ilvD* from PGAL10. We found that sorted strains produce an average of 296 ± 19 mg/L isobutanol from 15% galactose, with the highest producing strain (named YZy452) achieving 310 ± 15 mg/L isobutanol (**Supplementary Fig. 9**). In contrast, 20 randomly picked colonies from the unsorted population produce on average only 188 ± 19 mg/L, close to the 169 ± 12 mg/L produced in the baseline strain (YZy449) (**Supplementary Fig. 9**). This result demonstrates that our biosensor can also help identify strains with increased cytosolic isobutanol production, achieving titers as much as 83% higher than the baseline strain, and 65% higher than those of randomly picked colonies from the unsorted population.

We next used our biosensor to identify variants of *Ll_ilvD* that further enhance isobutanol production. Beginning with the highest producing strain isolated from the sorted population above (YZy452), we introduced an error-prone PCR library of *Ll_ilvD* mutants, each under the control of the constitutive PTDH3 promoter, in CEN plasmids. Thus, when the transformed strains are grown in glucose, the previously integrated copy of *Ll_ilvD* controlled by PGAL10 is repressed, and only the *Ll_ilvD* variant from the CEN plasmid library is expressed. After subjecting the *Ll_ilvD* library to three rounds of FACS, we isolated 24 random colonies from each round of sorting. The fraction of colonies producing more isobutanol than the control strain transformed with wild-type *Ll_ilvD* (YZy454) increases after each round of sorting (**Supplementary Fig. 10a**). By the third round of sorting, all 24 colonies examined produce between 150 mg/L and 260 mg/L isobutanol from 15% glucose, which is 2- to 3.5-times higher than the production observed with YZy454.

To confirm that the enhancement in isobutanol production in these sorted strains is caused by mutations in *Ll_ilvD*, we carried out fermentations with the sorted, cured, and retransformed strains as described above for Ilv6p and Leu4p variants, and measured their fluorescence intensity. We found six unique *Ll_ilvD* sequences among the 24 strains examined from the third round of sorting (**Supplementary Table 7**). Strains cured from plasmids containing these *Ll_ilvD* variants have undetectable levels of isobutanol production (**Fig. 5b**) and basal levels of GFP fluorescence intensity (**Supplementary Fig. 10b**), confirming that these variants are required for both phenotypes. Four out of the six retransformed strains reproduce the isobutanol titers of the sorted strains, including that retransformed with *Ll_ilvD* mutant #1, which retains 3.1-fold higher isobutanol production than the wild-type (**Fig. 5b**). The strain retransformed with *Ll_ilvD* mutant #2 produces less isobutanol than its sorted counterpart, suggesting that background mutations in the sorted strain contribute to its enhanced phenotype. However, despite its decrease in production relative to the originally sorted strain, the strain retransformed with *Ll_ilvD* mutant #2 still produces significantly more isobutanol than the strain overexpressing the wild-type *Ll_ilvD* (YZy454). Two of the other variants (mutants #1 and #3) also show statistically significant improvements in isobutanol titers relative to YZy454 (**Fig. 5b**).

We next mapped the mutations found in the three *Ll_ilvD* mutants that significantly improve isobutanol production onto the crystal structure of an IlvD homolog (**Supplementary Figure 11**). *Ll_*IlvD mutants #1 and #3 each have a single mutation, I433V and K535R, respectively, implying these residue substitutions are solely responsible for the enhancement in isobutanol production observed with each mutant. However, mutant #2 has three residue substitutions (V12A, S189P, and H439R), which makes it more difficult to ascertain the importance and contribution of each mutation. Nevertheless, the location of each mutation makes it possible to propose interesting hypotheses. All five mutations in the three variants are solvent exposed (**Supplementary Figure 11a, b**), suggesting that some of them may improve the solubility, expression, or stability of the enzyme. In addition, mutations I433V and S189P, from mutant #1 and mutant #2, respectively, are located on opposing lobes that define the putative substrate entrance to the active site (**Supplementary Figure 11c, d**). These lobes are known to undergo a significant conformational change^62^. In one conformation, the lobes seal the active site away from the solvent surrounding the enzyme (**Supplementary Figure 11c**), presumably to protect the 2Fe-2S cluster in the active site from oxidation. In the other conformation, the lobes move apart from each other to open the entrance to the active site, probably to allow substrates and products to enter and exit (**Supplementary Figure 11d**). This mechanism suggests that there is a balance between protecting the catalytic 2Fe-2S cluster from oxidation and allowing substrate and product exchange in the active site. Therefore, it is an intriguing possibility that mutations I433V and S189P improve enzyme activity by shifting this balance to one more optimal for the conditions in the yeast cytosol. Finally, mutation V12A, found in mutant #2, is located near the N-terminus of the enzyme, in a region involved in packing the tetramer and stabilizing the C-terminal H579, which coordinates the catalytic Mg^2+^ in the active site (**Supplementary Figure 11e**). It is possible that this mutation increases the stability of these structural features to enhance enzymatic activity. While careful biochemical characterization is essential to support or disprove these hypotheses, it is clear that our biosensor is capable of identifying new enzyme mutations that not only enhance isobutanol production, but also have the potential to further our molecular understanding of enzymatic mechanisms.

Cytosolic isobutanol production can be further enhanced by increasing the level of expression of *Ll*_*ilvD*. After retransforming the basal strain YZy452 (**Supplementary Table 1**) with a CEN plasmid containing a single copy of the best *ilvD* variant, *Ll*_*ilvD^I433V^*, the resulting strain (YZy469) produces 230 ± 13 mg/L of isobutanol from 15% glucose, which is 3.1-times higher than that of YZy454, harboring a single copy of the wild-type *Ll_ilvD* also in a CEN plasmid (**Fig. 5c**). When the same *Ll_ilvD^I433V^* mutant is introduced in a high copy 2µ plasmid, resulting in strain YZy470, the titer further increases to 334 mg/L ± 17 mg/L, 2.6-fold higher than overexpressing the wild-type *Ll_ilvD* in a 2µ plasmid (YZy468) (**Fig. 5c**). These results suggest that isobutanol production in the cytosol is limited by both *Ec_ilvC^P2D1-A1^* expression (YZy449 compared to YZy452, **Supplementary Fig. 9**) as well as *Ll_ilvD^I433V^* expression (YZy469 compared to YZy470, **Fig. 5c**), and that our biosensor is effective at increasing metabolic flux through both of these cytosolic bottlenecks.

Finally, we tested the ability of our biosensor to select for strains with enhanced metabolic flux through the cytosolic isobutanol biosynthetic pathway. We previously showed that optogenetically controlling the expression of the pyruvate decarboxylase encoded by *PDC1* and the mitochondrial isobutanol pathway in a triple *PDC* deletion background (*pdc1Δ pdc5Δ pdc6Δ*) boosts isobutanol production^63^. This is an effective way to increase metabolic flux towards pathways that compete with *PDC1*, which diverts pyruvate towards ethanol production, without completely eliminating pyruvate decarboxylase activity, which causes a severe growth defect in glucose^64, 65^. Thus, we tested our biosensor in a strain in which *PDC1* is controlled by *OptoEXP*^63^, an optogenetic circuit that induces expression only in blue light (450 nm), and the first enzyme of the cytosolic pathway (*Bs_*AlsS) is controlled by *OptoINVRT7*^66^, an optogenetic circuit that represses gene expression in the same wavelength of blue light and activates it in the dark (**Fig. 6a**). The starting strain for this experiment (YZy502) has a triple *PDC* deletion background with the dark-inducible cytosolic isobutanol pathway and light-inducible *PDC1* integrated into the *δ*-sites of a strain containing the isobutanol biosensor (**Supplementary Table 1**). As expected, this strain exhibits light-dependent growth on glucose, and its GFP fluorescence intensity and isobutanol production are higher in the dark than in the light, achieving as much as 296 mg/L ± 40 mg/L in dark fermentations from only 2% glucose (**Fig. 6b**). We transformed YZy502 with a 2µ plasmid (pYZ350, **Supplementary Table 2**) containing two copies of *Ec*_*ilvC^P2D1-A1^* and one copy each of *Ll*_*IlvD^I433V^*, *ARO10* (KDC), and *Ll_adhA^RE1^*(ADH)^67^. Using the isobutanol biosensor, we carried out two rounds of FACS from a library of pooled transformants to isolate YZy505, which produces 681 mg/L ± 29 mg/L isobutanol in dark fermentations from 2% glucose (**Fig. 6b**). This represents a yield of 34.1 ± 1.5 mg isobutanol per g glucose, which is comparable to previously reported cytosolic and mitochondrial isobutanol yields^24, 25, 32–34, 63^ (see Discussion). Thus, our isobutanol biosensor is equally adept at assisting in the construction of mitochondrial and cytosolic isobutanol pathways, and is effective at screening and isolating high-producing strains, including from optogenetically controlled strains with enhanced flux towards isobutanol production.

**Figure 6.**
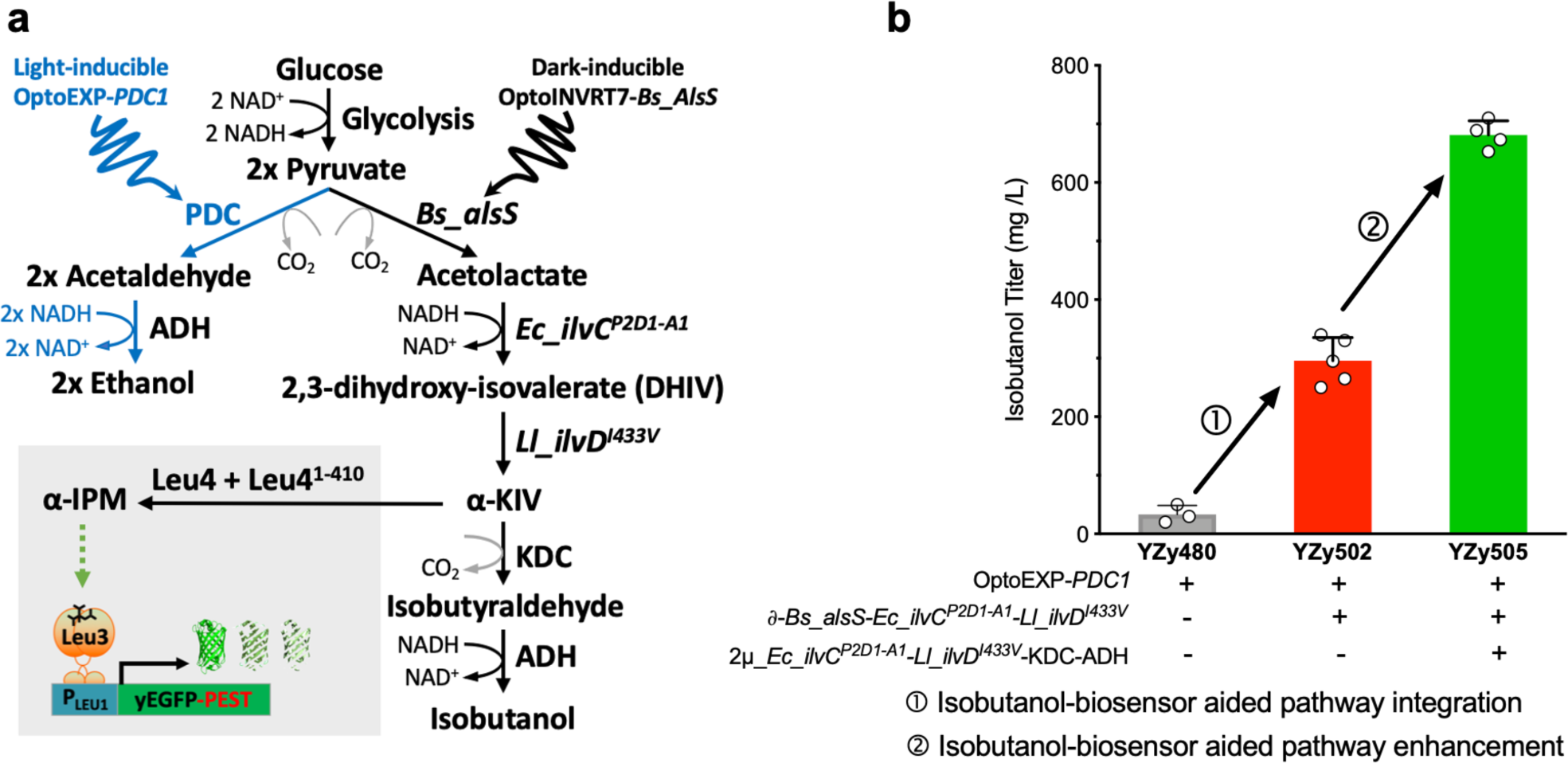
Biosensor-assisted engineering of a cytosolic isobutanol pathway containing optogenetic controls of *PDC1* and ALS for metabolic flux enhancement. (**a**) Schematic of the dark-inducible cytosolic isobutanol pathway (with *Bs_alsS* controlled by *OptoINVRT7*) and light-inducible *PDC1* (controlled by *OptoEXP*) in a triple *pdcΔ* strain containing the isobutanol biosensor. Isobutanol production of optogenetically regulated strains, including a basal strain containing only light-inducible *PDC1* (YZy480, gray); a strain also containing the dark-inducible upstream cytosolic pathway (*Bs_alsS, Ec_ilvC^P2D1-A1^,* and *Ll_ilvD^I433V^*) integrated in *δ*-sites (YZy502, red); and a strain containing additional enzymes from the cytosolic isobutanol pathway (*Ec_ilvC^P2D1-A1^, Ll_ilvD^I433V^*, KDC, and ADH) introduced in a 2µ plasmid (YZy505, green). Isobutanol production was measured after 48h fermentations in media containing 2% glucose. Strains YZy502 and YZy505 where each isolated after two rounds of FACS. Error bars represent the standard deviation of at least three biological replicates.

## Discussion

Genetically encoded biosensors have been developed to measure and control microbial biosynthesis of several products of interest^68–77^, including isobutanol in *E. coli*^42^. While *S. cerevisiae* is generally considered a preferred host for the industrial production of BCHAs, including isobutanol^78–80^, no genetically encoded biosensor for this class of chemicals has been reported in this organism. Here we show that by exploiting a transcriptional regulator of BCAA biosynthesis, Leu3p, it is possible to build biosensors specific to the BCHAs isobutanol and isopentanol.

Because Leu3p responds selectively to *α*-IPM, which is both a byproduct of isobutanol biosynthesis and an intermediate in isopentanol biosynthesis, our two biosensors are highly specific to each of these alcohols. Deleting *LEU2* makes the biosensor respond exclusively to isobutanol production (R^2^ = 0.97), as isopentanol biosynthesis from glucose is blocked in the absence of *LEU2*. By contrast, maintaining *LEU2* expression makes the biosensor response highly specific to isopentanol production (R^2^ = 0.98), but not to isobutanol (R^2^ < 0.20) (**Supplementary Fig. 12**). This high selectivity stems from the Leu3p mechanism of activation, which makes our biosensors substantially less likely to cross-react with other alcohols as compared to previously reported alcohol biosensors in *E. coli*^41, 42^ and *S. cerevisiae*^81^, which directly monitor alcohol concentrations, and can thus have significant cross-reactivity with several linear and branched chain alcohols^41, 42, 81^. Furthermore, by monitoring the concentration of an intracellular metabolite (*α*-IPM), as opposed to the concentration of the end products (isobutanol^42, 77^ or isopentanol), which could traverse cell membranes, our biosensors are better able to capture BCHA biosynthesis in each individual cell (using flow cytometry), rather than the average BCHA production in the fermentation. With our biosensor capabilities, we are able to screen and isolate high-producing strains from mixed populations, engineer metabolic enzymes involved in BCHA biosynthesis, and guide the construction and improvement of metabolic pathways for BCHA production in both mitochondria and the cytosol. Starting with random mutagenesis, and employing our biosensors and FACS, it is possible to identify enzyme variants with enhanced activity. With the assistance of our biosensors, we isolated several hyperactive mutants of three different proteins involved in BCHA production, including Ilv6p (**Fig. 3b**) and *Ll_*IlvD (**Fig. 5b**) for isobutanol production, and Leu4p (**Fig. 4c**) for isopentanol production. These enzymes are active in mitochondria (Ilv2p/Ilv6p), the cytosol (*Ll_*IlvD), or both (Leu4p), demonstrating that our biosensors can be used to engineer enzymes in either compartment, for the benefit of the mitochondrial or cytosolic isobutanol pathways, as well as the isopentanol pathway split across both compartments. For Ilv6p and Leu4p, the mutants we isolated enable the production of significantly more isobutanol and isopentanol, respectively, than strains harboring the wild-type enzymes or previously reported hyperactive mutants. To our knowledge, this is the first *Ll_*IlvD mutant reported to enhance isobutanol production. It may be possible to find even more active mutants by enriching the diversity of our libraries. However, this approach is ultimately limited by the ability of random mutations to further improve enzyme activity. It is also limited by the extent to which the catalytic role of each enzyme remains a bottleneck in the metabolic pathway, which determines whether enhancing the enzyme’s activity results in increased BCHA production, and thus biosensor signal. Nevertheless, the three examples we explored show that our biosensors are effective at finding hyperactive mutants of mitochondrial and cytosolic enzymes, and suggest that other metabolic enzymes and regulatory proteins involved in isobutanol and/or isopentanol production could be improved using this approach.

In addition to being instrumental for engineering individual enzymes, our biosensors can assist in the construction and improvement of entire metabolic pathways in either mitochondria or the cytosol, using two different approaches. For the mitochondrial isobutanol pathway, we used the biosensor to identify the highest producers from a library of strains containing random combinations of the pathway genes, all encoding enzymes targeted to mitochondria. Using the biosensor allowed us to find strains that produce, on average, more than twice as much isobutanol as strains randomly selected without the assistance of our biosensor (**Fig. 2**). For the cytosolic pathway, we first used our biosensor to open specific metabolic bottlenecks, finding strains with higher copy number of *Ec_ilvC^P2D1-A1^* (**Supplementary Fig. 9**). We then employed the biosensor to identify the hyperactive *Ll*_*IlvD^I433V^* mutant, thereby enhancing isobutanol production from glucose (**Fig. 5c**). Finally, we utilized the biosensor to isolate the highest producers from a library of optogenetically controlled strains with increased metabolic flux through the cytosolic pathway (**Fig. 6b**). Therefore, our isobutanol biosensor is effective in assisting with the construction and improvement of both mitochondrial and cytosolic isobutanol pathways, using a variety of strategies to enhance production. Previous efforts to construct cytosolic isobutanol pathways have involved decompartmentalizing the ALS, KARI, and DHAD enzymes encoded by the endogenous *ILV* genes by deleting their mitochondrial localization signals^32–34^. By contrast, the cytosolic isobutanol pathway we systematically built with our isobutanol biosensor uses heterologous bacterial genes to express these enzymes in the cytosol. The best strain we obtained (YZy505) produces 681 mg/L ± 29 mg/L of isobutanol with a yield of 34.1 ± 1.5 mg isobutanol per g glucose, comparable to similarly engineered cytosolic strains^33, 34, 82^. This production could be improved by incorporating additional deletions previously reported to eliminate competing pathways^33^ or by increasing metabolic flux through the isobutanol pathway with the use of light pulses during the production phase of fermentations with our optogenetically controlled strain^63^. Moreover, our biosensor is well positioned to accelerate the discovery of new gene deletions and optimal light schedules to further improve isobutanol production.

Our biosensors are also applicable to several lines of research in biotechnology, basic science, and metabolic control. Their most immediate application will likely be to accelerate the development of strains for the production of isobutanol and isopentanol as promising advanced biofuels. However, because these biosensors are controlled by the activity of the BCAA regulator Leu3p, rather than directly sensing isobutanol or isopentanol concentrations, they may also be useful to develop strains for the production of valine, leucine, or other products derived from their metabolism^9–16^. An advantage of biosensors based on transcription factors is that they can be used to control the expression of any gene of interest. Therefore, Leu3p can be used to control the expression of not only reporter genes to develop high-throughput screens for strains with enhanced production (as demonstrated in this study), but also of essential genes to develop high-throughput selection assays^41, 83^, or of key metabolic genes to develop autoregulatory mechanisms for dynamic control of product biosynthesis^43, 84^. For the same reason (that Leu3p regulates BCAA biosynthesis, which initiates in mitochondria) these biosensors are potentially useful to study fundamental questions in BCAA metabolism or mitochondrial activity. The two previously described BCAA biosensors^85, 86^ measure collective intracellular levels of valine, leucine, and isoleucine, limiting their applicability. In addition, while biosensors based on fluorescence or bioluminescence resonance energy transfer, or GFP-ligand-binding-protein chimeras have been developed to probe the mitochondrial metabolic state^87^, our biosensor is, to the best of our knowledge, the first mitochondrial biosensor based on a transcription factor, and the only one to probe the flux through a mitochondrial metabolic pathway. Finally, the fact that the isobutanol biosensor is functional in strains carrying optogenetic controls for the production of this biofuel raises the intriguing possibility of simultaneously monitoring and controlling isobutanol production, which may offer new research avenues in closed loop metabolic control and cybergenetics for metabolic engineering.

## Methods

### Chemicals, reagents and general molecular biology techniques

All chemicals and solvents were obtained from Sigma-Aldrich (St. Louis, MO, USA). Phusion High-Fidelity DNA Polymerase, Taq DNA polymerase, T4 DNA ligase, T5 Exonuclease, Taq DNA ligase, Calf Intestinal Alkaline Phosphatase (CIP), Deoxynucleotide (dNTP) solution mix, and restriction enzymes were purchased from New England BioLabs (NEB, Ipswich, MA, USA) or Thermo Fisher Scientific (Waltham, MA, USA). QIAprep Spin Miniprep, QIAquick PCR Purification, and QIAquick Gel Extraction Kits (Qiagen, Valencia, CA, USA) were used for plasmid isolation and DNA fragments purification according to the manufacturer’s protocols. Genomic DNA extractions were carried out using the standard phenol chloroform procedure^88^. Genotyping PCR assays were performed using GoTaq Green Master Mix (Promega, Madison, WI, USA). The oligonucleotides used in this study (**Supplementary Table 8**) were obtained from Integrated DNA Technologies (IDT, Coralville, IA, USA). *Escherichia coli* DH5α was used for routine transformations. All of the constructed plasmids were verified by DNA sequencing (GENEWIZ, South Plainfield, NJ, USA).

### Growth media

Unless otherwise specified, yeast cells were grown at 30 °C on either YPD medium (10 g/L yeast extract, 20 g/L peptone, 0.15 g/L tryptophan and 20 g/L glucose) or synthetic complete (SC) drop out medium (20 g/L glucose, 1.5 g/L yeast nitrogen base without amino acids or ammonium sulfate, 5 g/L ammonium sulfate, 36 mg/L inositol, and 2 g/L amino acid drop out mixture) supplemented with 2% glucose, or non-fermentable carbon sources such as 2% galactose, or the mixture of 3% glycerol and 2.5% ethanol. 2% Bacto™-agar (BD, Franklin Lakes, NJ, USA) was added to make agar plates.

### Assembly of DNA constructs

DNA construction was performed using standard restriction-enzyme digestion and ligation cloning and isothermal assembly (a.k.a. Gibson Assembly)^89^. Endogenous *S. cerevisiae* genes (*ILV6*, *LEU4*, and *AFT1*) were amplified from genomic DNA of CEN.PK2-1C by polymerase chain reaction (PCR) using a forward primer containing an NheI restriction recognition site and a reverse primer containing an XhoI restriction recognition site (**Supplementary Table 8)**. This enabled subcloning of PCR-amplified genes into pJLA vectors^24^ and pJLA vector-compatible plasmids. Genes from other organisms, including *Bs_alsS*, *Ec*_*ilvC^P2D1-A1^* and *Ll_ilvD* were codon optimized for *S. cerevisiae* and synthesized by Bio Basic Inc. (Amherst, NY, USA). These genes were designed with flanking NheI and XhoI sites at the 5’ and 3’ ends, respectively, and without restriction sites for XmaI, MreI, AscI, PmeI, PacI, and other relevant restriction enzymes. All plasmids constructed or used in this study are listed in **Supplementary Table 2**.

### Constructing an isobutanol biosensor

The reporter for the isobutanol biosensor was constructed by placing the *S. cerevisiae LEU1* promoter (PLEU1), in front of the yeast-enhanced green fluorescent protein (yEGFP) fused to a PEST tag. The *LEU1* promoter was amplified from genomic DNA of CEN.PK2-1C using primers Yfz_Oli31 and Yfz_Oli32. The yEGFP-PEST fragment was amplified from plasmid pJA248 using primers Yfz_Oli33 and Yfz_Oli34. These two PCR fragments were inserted by Gibson assembly into the pJLA vector JLAb131 to make an intermediate plasmid (pYZ13) containing the fragment PLEU1-yEGFP_PEST-TADH1, which was then subcloned into a *HIS3* locus integration (His3INT) vector pYZ12B^63^, yielding the intermediate plasmid pYZ14. To overcome the leucine inhibition of Leu4p, we used a Leu4 mutant lacking the regulatory domain (Leu4^1-410^). The truncated *LEU4^1-410^* was amplified from genomic DNA of CEN.PK2-1C using primers Yfz_Oli35 and Yfz_Oli36 and subcloned into a pJLA vector plasmid JLAb23, which contains the constitutive promoter PTPI1 and terminator TPGK1. The PTPI1-*LEU4*^1-410^-TPGK1 cassette from the resulting plasmid pYZ2 was then subcloned into pYZ14 using sequential gene insertion cloning^24^ to form the plasmid pYZ16 (**Supplementary Table 2**).

### Constructing an isopentanol biosensor

To construct the isopentanol biosensor, we removed the PEST-tag of the yEGFP reporter in plasmid pYZ14 by annealed oligo cloning. The two single-stranded overlapping oligonucleotides Yfz_Oli59 and Yfz_Oli60 were annealed and cloned directly into the overhangs generated by restriction digest of pYZ14 at SalI and BsrGI sites. The resulting plasmid (pYZ24) contains cassette PLEU1-yEGFP-TADH1. Next, we added a catalytically active and leucine-insensitive *LEU4* mutant (*LEU4^ΔS547^*, a deletion in Ser547)^47,48^ to pYZ24 to form the isopentanol biosensor pYZ25. The deletion in Ser547 was achieved by site-directed mutagenesis using a plasmid containing wild type *LEU4* (*LEU4^WT^*) amplified from genomic DNA of CEN.PK2-1C as the template. *LEU4^ΔS547^* was then subcloned into pJLA vector JLAb23 to form plasmid pYZ1. The PTPI1-*LEU4^ΔS547^*-TPGK1 cassette from pYZ1 was then subcloned into pYZ24 to form the plasmid pYZ25.

### Constructing template plasmids for error-prone PCR

The backbone vector pYZ125, used to prepare the random mutagenesis libraries, was constructed by subcloning a DNA fragment containing a *TDH3* promoter (PTDH3), a KOZAK sequence, a multiple cloning site (MCS) sequence flanked by NheI and XhoI sites, a stop codon, and an ADH1 terminator (TADH1) from plasmid JLAb131 into the CEN plasmid pRS416^90^. *ILV6*, *LEU4,* and *Ll*_*ilvD* were subcloned into pYZ125, after linearization with NheI and XhoI, to create pYZ127 (harboring *ILV6^WT^)*, pYZ149 (harboring *LEU4^WT^*), and pYZ126 (harboring *Ll_ilvD^WT^*), respectively. The *ILV6^V90D/L91F^* mutant was made using QuikChange site-directed mutagenesis and subcloned into pYZ125 to create pYZ148 (harboring *ILV6^V90D/L91F^* mutant)^25^. Leucine inhibition insensitive *LEU4* mutants *LEU4^ΔS547^* and *LEU4^1-410^* were subcloned into pYZ125 from pYZ1 and pYZ2, respectively, to form pYZ154 and pYZ155 (**Supplementary Table 2**).

### Constructing δ-integration cassettes

We used a previously developed *δ*-integration (*δ*-INT) vector, pYZ23^63^, to integrate multiple copies of gene cassettes into genomic YARCdelta5 δ-sites, the 337 bp long-terminal-repeat of *S. cerevisiae* Ty1 retrotransposons (YARCTy1-1, SGD ID: S000006792). The selection marker in pYZ23 is the shBleMX6 gene, which encodes a protein conferring resistance to zeocin and allows selection of different numbers of integration events by varying zeocin concentrations^91, 92^. Resistance to higher concentrations of zeocin correlates with a higher number of gene cassettes integrated into δ-sites. The δ-integration plasmids pYZ33, pYZ113, and pYZ34 were constructed by subcloning gene cassettes from plasmids pJA123, JLAb581, and pJA182, respectively. We used restriction site pairs XmaI/AscI (to extract gene cassettes) and MreI/AscI (to open pYZ23)^24^. Plasmid pYZ33 contains the

*δ*-integration cassette of the upstream isobutanol synthetic pathway *ILV* genes (*δ*-INT-*ILV*s); plasmid pYZ113 contains the *δ*-integration cassette of the mitochondria-targeted downstream Ehrlich pathway (*δ*-INT-CoxIVMLS-*ARO10*, CoxIVMLS-*Ll_adhA^RE1^*); plasmid pYZ34 contains the *δ*-integration cassette of the full mitochondria-targeted isobutanol synthetic pathway (*δ*-INT-ILVs, CoxIVMLS-*ARO10*, CoxIVMLS-*Ll_adhA^RE1^*). Plasmid pYZ417 contains the dark-inducible cytosolic isobutanol pathway and the light-inducible *PDC1* (*δ*-integration-OptoEXP-*PDC1*-OptoINVRT7-cytosolic isobutanol pathway). It was constructed by sequentially inserting cassettes PTDH3-*Ec_ilvC^P2D1-A1^*-TCYC1 (from pYZ196), PTEF1-*Ll_ilvD^I433V^*-TTPS1 (from pYZ383), and PGal1-S-*Bs_alsS*-TACT1 (from pYZ384) into the *δ*-integration plasmid EZ-L235^63^, which contains the OptoEXP-*PDC1* cassette (PC120-*PDC1*-TACT1). All of the *δ*-integration plasmids were linearized with PmeI prior to yeast transformation.

### Error-prone PCR and random mutagenesis library construction

The random mutagenesis libraries of *ILV6*, *LEU4* and *Ll_ilvD* were generated by error-prone PCR using the GeneMorph II Random Mutagenesis Kit (Agilent Technologies, Santa Clara, CA, USA). The CEN plasmids harboring wild-type *ILV6* (pYZ127), *LEU4* (pYZ149), *Ll_ilvD* (pYZ126) or the *ILV6^V90D/L91F^* mutant (pYZ148) were used as templates for error-prone PCR. Various amounts of DNA template were used in the amplification reactions to obtain low (0-4.5 mutations/kb), medium (4.5-9 mutations/kb), and high (9-16 mutations/kb) mutation frequencies as described in the product manual. Primers Yfz_Oli198 and Yfz_Oli242, which contain NheI and XhoI restriction sites at their 5’ ends, respectively, were used for amplifying and introducing random mutations to the region between the start codon and stop codon of each gene. The PCR fragments were incubated with DpnI for 2 h at 37°C to degrade the template plasmids before being purified using a PCR purification kit (Qiagen). The purified PCR fragments were digested overnight at 37°C using NheI and XhoI and ligated overnight at 16°C to backbone vector pYZ125, opened with NheI and XhoI. To create an ep_library of the leucine regulatory domain of Leu4p, the forward primer Yfz_Oli243, containing a BglII restriction site at its 5’ end, was used together with the reverse primer Yfz_Oli242. The purified PCR fragments were digested overnight at 37°C using BglII and XhoI and ligated overnight at 16°C to backbone vector pYZ149, opened with BglII and XhoI. The plasmid libraries that resulted were transformed into ultra-competent *E. coli* DH5α and plated onto LB-agar plates (five 150 mm petri dishes per library) containing 100 µg/ml of ampicillin. After incubating the plates overnight at 37°C, the resulting lawns were scraped off the agar plates and the plasmid libraries were extracted using a QIAprep Spin Miniprep Kit (QIAGEN). The plasmid libraries were subsequently used for yeast transformation.

### Yeast strains and yeast transformations

*Saccharomyces cerevisiae* strain CEN.PK2-1C (*MAT**a** ura3-52 trp1-289 leu2-3,112 his3-1 MAL2-8c SUC2*) and its derivatives were used in this study (**Supplementary Table 1**). Deletions of *BAT1*, *BAT2*, *ILV6*, *LEU4*, *LEU9*, *ILV3, TMA29, PDC1*, *PDC5*, *PDC6*, and *GAL80* were obtained using PCR-based homologous recombination. DNA fragments containing lox71- and lox66-flanked antibiotic resistance cassettes were amplified with PCR from pYZ55 (containing the hygromycin resistance gene hphMX4), pYZ17 (containing the G418 resistance gene KanMX), or pYZ84 (containing the nourseothricin resistance gene NAT1), using primers with 50-70 base pairs of homology to regions upstream and downstream of the ORF of the gene targeted for deletion^49^. Transformation of the gel-purified PCR fragments was performed using the lithium acetate method^93^.

Cells transformed using antibiotic resistance markers, were first plated onto nonselective YPD plates for overnight growth, then replica plated onto YPD plates containing 300 µg/mL hygromycin (Invitrogen, Carlsbad, CA), 200 µg/mL nourseothricin (WERNER BioAgents, Jena, Germany), or 200 µg/mL Geneticin (G-418 sulfate) (Gibco, Life Technologies, Grand Island, NY, USA). Chromosomal integrations and plasmid transformations were performed using the lithium acetate method^93^. Cells transformed using auxotrophic markers were plated onto agar plates containing corresponding synthetic complete (SC) drop out medium. Cells transformed for *δ*-integration were first incubated in YPD liquid medium for six hours and then plated onto nonselective YPD agar plates for overnight growth. The next day, cells were replica plated onto YPD agar plates containing 200 µg/mL (for FACS), or higher concentration (500, 800, 1200, and 1500 µg/mL for titrating higher copy numbers of integrations) of Zeocin (Invitrogen, Carlsbad, CA), and incubated at 30°C until colonies appeared. Cells transformed with plasmid libraries or 2µ plasmids (used for FACS) were plated on multiple large agar plates (three 150 mm petri dishes per library). The resulting lawns were scraped off the agar plates and used for flow cytometry assays and FACS (see below). All strains with gene deletions or chromosomal integrations (expect the *δ*-integration) were genotyped with positive and negative controls to confirm the removal of the ORF of interest or the presence of integrated DNA cassette.

### Strains to obtain correlations between biosensor signals and BCHA production

To construct strains used to obtain the correlation between the isobutanol biosensor fluorescence intensity and isobutanol production, we first integrated the isobutanol biosensor into the *HIS3* locus of either a wild-type CEN.PK2-1C strain or a *bat1*Δ strain (SHy1). The resulting strains (YZy121 and YZy81) were then transformed with a linearized *δ*-integration cassette containing either the *ILV* genes alone (pYZ33), or the *ILV* genes combined with KDC, and ADH targeted to the mitochondria (pYZ34). Seven strains with different isobutanol production levels were selected using increasing Zeocin (500, 800, 1200, or 1500 µg/mL). To construct strains used to measure the correlation between the isopentanol biosensor fluorescence intensity and isopentanol production, we first restored the *LEU2* gene and deleted *BAT1, LEU4*, and *LEU9* in the wild-type CEN.PK2-1C strain, resulting in strain YZy140. We then integrated the isopentanol biosensor (pYZ25) into the *HIS3* locus to yield strain SHy134. We also constructed the strain SHy181, which has the native *BAT1* restored into the *TRP1* locus. Finally, six strains with different isopentanol production capabilities were constructed by transforming SHy134 and SHy181 with CEN/ARS vectors harboring *ILV1* (JLAb705), a cassette containing *ILV2*, *ILV3*, and *ILV5* (JLAb691), or empty vector (pYZ125).

### Strains with cytosolic isobutanol pathway

We first constructed a baseline strain for cytosolic isobutanol production (YZy449). We used constitutive promoters to express *Bs_alsS*, *Ec_ilvC^P2D1-A1^,* and *AFT1*, and the galactose-inducible and glucose-repressible promoter PGAL10 to express *Ll_ilvD*. This approach allows us to use the same strain to screen for *Ec_ilvC^P2D1-A1^* and *Ll_ilvD* mutants using different carbon sources (galactose and glucose, respectively). We first deleted *ILV3* and *TMA29* in strain YZy121, which contains the isobutanol biosensor, resulting in strain YZy443. We then transformed strain YZy443 with the *URA3* integration cassette from plasmid pYZ196 (loxP-*URA3*-loxP-PTDH3-*AFT1*-TADH1-[PTEF1-*Bs_alsS*-TACT1-PTDH3-*Ec_ilvC^P2D1-A1^*-TADH1]-P_GAL10_-*Ll_ilvD*-TACT1), resulting in strain YZy447. In order to use the *URA3* marker in later experiments, we recycled the *URA3* marker in YZy447 using the Cre-loxP site-specific recombination system^94^ (using plasmid pSH63) and counter selected in YPD plates with 1 mg/mL of 5-FOA (see below), resulting in the final strain YZy449. Both YZy447 and YZy449 make 169 mg/L (169 ± 10 mg/L and 169 ± 12 mg/L, respectively) isobutanol in fermentations using galactose as carbon source but are unable to make isobutanol from glucose (**Supplementary Fig. 13**).

### Isobutanol biosensor in optogenetically controlled strains with enhanced flux through the cytosolic isobutanol pathway

To combine the isobutanol biosensor with optogenetic controls of the cytosolic isobutanol pathway, we integrated OptoINVRT7^66^ (using EZ-L439) into the *HIS3* locus of YZy90, a *gal80Δ*, triple *pdcΔ* (*pdc1Δ pdc5Δ pdc6Δ*) strain containing a constitutively expressed copy of *PDC1* in a 2μ plasmid, resulting in YZy480. We removed the 2μ-*PDC1* plasmid using 5-FOA (see below). YZy480 was first grown in SC medium supplemented with 3% glycerol and 2.5% ethanol (SCGE) overnight and then streaked on an SCGE agar plate containing 1 mg/mL 5-FOA (see below). The resulting strain (YZy481) is able to grow in medium supplemented with glycerol and ethanol, but not in medium supplemented with glucose. We next introduced the isobutanol biosensor into the *GAL80* locus of YZy481 using the cassette from plasmid pYZ414. We measured GFP fluorescence intensity to confirm the isobutanol biosensor was integrated successfully. The GFP-positive strain, YZy487, was confirmed by genotyping PCRs. A dark-inducible cytosolic isobutanol pathway and the light-inducible *PDC1* were then introduced to strain YZy487 via *δ*-integration using linearized pYZ417. After two rounds of FACS, we analyzed the GFP fluorescence and isobutanol titers of ten colonies in dark fermentations with 2% glucose (see below). The colony with the highest GFP fluorescence intensity, corresponding to the highest isobutanol titer, was chosen as the host strain (YZy502) for further enhancement of the cytosolic isobutanol production. To achieve a strain with even higher metabolic flux through the cytosolic isobutanol pathway, we introduced a 2µ plasmid pYZ350, containing partial cytosolic isobutanol pathway genes, and applied FACS to isolate high-producing transformants.

### Flow cytometry/FACS

A BD LSRII Multi-Laser flow cytometer equipped with FACSDiva software V.8.0.2 (BD Biosciences, San Jose, CA) was used to quantify yEGFP fluorescence at an excitation wavelength of 488 nm and an emission wavelength of 510 nm (525/50 nm bandpass filter). Cells were gated on forward scatter (FSC) and side scatter (SSC) signals to discard debris and probable cell aggregates. The typical sample size was 50,000 events per measurement. The median fluorescence intensity (MFI) of the gated cell population was calculated by the software. A BD FACSAria Fusion flow cytometer with FACSDiva software was used for fluorescence activated cell sorting (FACS) with a 488 nm excitation wavelength and a bandpass filter of 530/30 for yEGFP detection. Cells exhibiting high levels of GFP fluorescence (top ∼1%) were sorted. Sorted cells were collected into 1 mL of the same medium (see below). 50 µL of each sample of collected sorted cells (∼1 mL) were streaked on a corresponding agar plate and incubated at 30°C to obtain 24 random colonies to evaluate the effectiveness of sorting. For each of the single colonies isolated, MFI and BCHA production were measured. The rest of the sorted cells (∼950 µL) were incubated at 30°C and at 200 rpm agitation to reach the stationary growth phase, followed by subculturing (1:100 dilution) in the same medium for the next round of sorting. FlowJo X software (BD Biosciences, San Jose, CA) was used to analyze the flow cytometry data.

### Sample preparation for flow cytometry assays and FACS high-throughput screens

Fluorescence measurements and FACS were performed on samples in mid-exponential growth phase. Single colonies from agar plates or yeast transformation libraries diluted 1:100 were first cultured overnight until stationary phase in synthetic complete (SC), or synthetic complete minus uracil (SC-ura) medium, supplemented with 2% glucose or galactose (for screening the best producers from a library of strains containing additional copies of *Ec_ilvC^P2D1-A1^*). Overnight cultures were diluted 1:100 in the same fresh medium and grown to mid-exponential growth phase (12-13 h after inoculation). SC medium supplemented with 2% glucose or galactose (for screening the library of strains containing additional copies of *Ec_ilvC^P2D1-A1^*) was used for flow cytometry assays and FACS of *δ*-integration libraries. SC-ura medium supplemented with 2% glucose was used for flow cytometry assays and FACS of strains containing plasmid libraries. The growth media used for flow cytometry assays and FACS of *ILV6*-samples contain four times more valine (2.4 mM) than the synthetic defined medium described above.

Strains with *PDC1* and *Bs_alsS* controlled by optogenetic circuits (*OptoEXP*^63^ and *OptoINVRT7*^66^, respectively) grow slower in medium supplemented with 2% glucose and produce isobutanol in the dark. We modified the growth conditions for these strains such that after inoculation with cells in stationary phase, cultures were incubated for 8 h under constant blue light, followed by 10 h of incubation in the dark (see below) before flow cytometry or FACS.

For all strains, cultures used for flow cytometry assays were diluted to an OD600 of approximately 0.1 with PBS. Samples used for FACS were diluted in fresh medium identical to their growth medium to OD600 of approximately 0.8.

### Measurement of the response of the biosensor to α-IPM and α-KIV

Measurements of the responses of the biosensor to α-IPM and α-KIV were carried out in a CEN.PK2-1C background strain, with the isobutanol biosensor integrated at the *HIS3* locus (YZy121). A single colony from an agar plate was cultured overnight in SC medium supplemented with 2% glucose. The overnight culture was diluted 1:100 in fresh SC medium supplemented with 2% glucose, and 1 mL of the diluted culture was added to each well of a 24-well cell culture plate (Cat. 229524, CELLTREAT Scientific Products, Pepperell, MA, USA). Then, 10 µL of freshly made α-IPM (pH 7.0) or α-KIV (pH 7.0) solutions were added to each well to reach different final concentrations ranging from 0 µM to 75 µM. The 24-well plate was shaken for 12 h in an orbital shaker (Eppendorf, New Brunswick, USA) at 30°C and at 200 rpm agitation. The GFP fluorescence of each sample was measured using flow cytometry when the cultures were in exponential phase.

### Yeast fermentations for isobutanol or isopentanol production

High-cell-density fermentations were carried out in sterile 24-well microtiter plates (Cat. 229524, CELLTREAT Scientific Products, Pepperell, MA, USA) in an orbital shaker (Eppendorf, New Brunswick, USA) at 30°C and at 200 rpm agitation. Single colonies were grown overnight in 1 mL of synthetic complete (SC), or synthetic complete minus uracil (SC-ura) medium, supplemented with 2% glucose. The next day, 10 µL of the overnight culture were used to inoculate 1 mL of SC (or SC-ura) medium supplemented with 2% glucose in a new 24-well plate. After 20 h, the plates were centrifuged at 1000 rpm for 5 min, the supernatant was discarded, and cells were re-suspended in 1 mL of SC (or SC-ura) supplemented with 15% glucose (or galactose). The plates were covered with sterile adhesive SealPlate^®^ films (Cat. # STR-SEAL-PLT; Excel Scientific, Victorville, CA) and incubated for 48 h at 30°C with 200 rpm shaking. The SealPlate^®^ films were used in all 24-well plate fermentations to maintain semi-aerobic conditions in each well, and to prevent evaporation and cross-contamination between wells. At the end of the fermentations, the OD600 of the culture in each well was measured in a TECAN infinite M200PRO plate reader (Tecan Group Ltd., Männedorf, Switzerland). Plates were then centrifuged for 5 min at 1000 rpm. The supernatant (approximately 1 mL) from each well was collected and analyzed using HPLC as described below.

Fermentations of strains with optogenetic controls were also carried out in sterile 24-well microtiter plates, at high-cell-density as described above, but with some modifications as previously described^63^. Blue LED panels (HQRP New Square 12” Grow Light Blue LED 14W, Amazon) were placed above the 24-well microtiter plates to stimulate cell growth with blue light at an intensity range of 70-90 µmol m^-2^ s^-1^ and duty cycles of 15 s on and 65 s off. The vertical distance between the LED panel and the top of the 24-well plates was adjusted based on the light intensity of each LED panel, measured with a spectrum quantum meter (Cat. MQ-510, Apogee Instruments, Inc., UT, USA). The light duty cycles were achieved by regulating the LED panels using a Nearpow Multifunctional Infinite Loop Programmable Plug-in Digital Timer Switch (purchased from Amazon). Single colonies were grown overnight in 1 mL of SC or SC-ura medium supplemented with 2% glucose in individual wells of the 24-well microtiter plates under constant blue light in an orbital shaker (Eppendorf, New Brunswick, USA) at 30°C and at 200 rpm agitation. The next day, each overnight culture was used to inoculate 1 mL of the same medium to reach an initial OD600 of 0.1 and grown at 30°C, 200 rpm, and under pulsed blue light (15 s on and 65 s off). We incubated the cultures in the light until they reached different cell densities (ρ), at which we switched them from light to dark. The 24-well plates were kept in the dark by wrapping them with aluminum-foil (and continuing to incubate at 30 °C, 200 rpm) for θ hours (the incubation time in the dark). After the dark incubation period, the cells were centrifuged and re-suspended in 1 mL of fresh SC-ura medium supplemented with 2% glucose. The plates were covered with sterile adhesive SealPlate^®^ films (Cat. # STR-SEAL-PLT; Excel Scientific, Victorville, CA) and incubated in the dark (wrapped in aluminum foil) for 48 h at 30°C and 200 rpm shaking. Subsequently, the cultures were centrifuged for 5 min at 1000 rpm, and the supernatants were collected and used for HPLC analysis.

The parameters ρ and θ used in the initial screening fermentations are an OD600 of 6 and 6 hours, respectively, which were estimated based on the characterization and modeling of the optogenetic circuits OptoEXP^63^ and INVRT7^66^. The optimal ρ (OD600 of 5) and θ (6 hours) of the best isobutanol-producing strains (YZy502 and YZy505) were determined experimentally by measuring the isobutanol titers from fermentations using different ρ and θ values. To vary these parameters, a single colony of each strain was first grown to stationary phase in SC (for YZy502) or SC-ura (for YZy505) medium supplemented with 2% glucose under constant blue light at 30°C. The overnight cultures were diluted to an OD600 of 0.1 in the same medium. We began incubations (at 30°C and 200 rpm) of the diluted cultures at different times and under pulsed blue light to achieve cultures with different OD600 values, which correspond to variations in ρ, ranging from 1 to 9. After measuring the OD600 of each culture, we switched them from light to dark and incubated for 6 hours at 30°C and 200 rpm. After the dark incubation period, the cells were centrifuged and re-suspended in 1 mL of the same medium supplemented with 2% glucose. The plates were covered with sterile adhesive SealPlate^®^ films (Cat. # STR-SEAL-PLT; Excel Scientific, Victorville, CA) and incubated in the dark (wrapped in aluminum foil) for 48 h at 30°C and 200 rpm. Subsequently, the cultures were collected and processed for HPLC analysis (see below). To determine the optimal θ, we diluted the overnight culture to an OD600 of 0.1 and incubated it at 30°C and 200 rpm, and under pulsed blue light to reach an OD600 of 5, which is the optimal θ determined in the previous experiment. Next, we incubated the cells in the dark (30°C and 200 rpm) for different numbers of hours, ranging from 1 to 10, which correspond to variations in θ. After the dark incubation period, cells were centrifuged and re-suspended in 1 mL of the same medium supplemented with 2% glucose, followed by 48 h of incubation in the dark (30°C and 200 rpm), sample collection, and HPLC analysis as described below.

### Removal of plasmids from *Saccharomyces cerevisiae*

*URA3* plasmids in yeast strains were removed by 5-fluoroorotic acid (5-FOA) selection^95^. Cells were first grown in YPD overnight and then streaked on SC agar plates containing 1 mg/mL 5-FOA (Zymo Research, Orange, CA, USA). A single colony from an SC/5-FOA agar plate was streaked again on a new SC/5-FOA agar plate. To confirm that strains were cured of the *URA3*-containing plasmid, they were inoculated into SC-ura medium supplemented with 2% glucose to test their growth phenotype. Strains in which the plasmid was removed were not able to grow in SC-ura medium.

### Yeast plasmid isolation

Plasmid isolation from yeast was performed according to a user-developed protocol from Michael Jones (protocol PR04, Isolation of plasmid DNA from yeast, QIAGEN) using a QIAprep Spin Miniprep kit (QIAGEN, Valencia, CA, USA)^96^. The isolated plasmids were retransformed into *E. coli* DH5α to produce plasmids at higher titers for subsequent sequencing and retransformation of the parental yeast strain.

### Analysis of chemical concentrations

The concentrations of glucose, isobutanol, and isopentanol were determined with high-performance liquid chromatography (HPLC) using an Agilent 1260 Infinity instrument (Agilent Technologies, Santa Clara, CA, USA). Samples were centrifuged at 13,300 rpm for 40 min at 4°C to remove residual cells and other solid debris, and analyzed using an Aminex HPX-87H ion-exchange column (Bio-Rad, Hercules, CA USA). The column was eluted with a mobile phase of 5 mM sulfuric acid at 55°C and with a flow rate of 0.6 mL/min for 50 min. The chemical concentrations were monitored with a refractive index detector (RID) and quantified by comparing the peak areas to those of prepared standards,

### Statistical analysis

Two-tailed Student’s *t*-tests were used to determine the statistical significance of differences observed in product titers between strains. Probabilities (*P*-values) less than (or equal to) 0.05 are considered sufficient to reject the null hypothesis (that the means of the two samples are the same) and accept the alternative hypothesis (that the means of the two samples are different).

## Acknowledgments

We thank Christina DeCoste and Katherine Rittenbach, who assisted with all flow cytometry instrumentation. This work was supported by the U.S. Department of Energy, Office of Science, Office of Biological and Environmental Research, Genomic Science Program under award number DE-SC0019363, as well as by the National Science Foundation, and NSF CAREER Award CBET-1751840 (to J.L.A.). This work was also supported by the NSF Graduate Research Fellowship Program grant DGE-1656466, the P.E.O. Scholar Award, and the Harold W. Dodds Fellowship from Princeton University (to S.K.H.). J.L.A. is also supported by The Pew Charitable Trusts, The Camille Dreyfus Teacher-Scholar Award, The Eric and Wendy Schmidt Transformative Technology Fund, and grants from Princeton University and the Andlinger Center for Energy and the Environment.

