## Supplemental Information for "Genetically encoded biosensors for branched-chain amino acid metabolism to monitor mitochondrial and cytosolic production of isobutanol and isopentanol in yeast"

#### Supplementary Tables

**Supplementary Table 1. Yeast strains used in this study.**

| Strain | Description | Genotype (Plasmid contents in parenthesis) | Source |
|---|---|---|---|
| CEN.PK2-1C | Wild-type *Saccharomyces cerevisiae* | *MAT**a** ura3-52 trp1-289 leu2-3,112 his3-1* $MAL2\text{-}8^{c}$ *SUC2* | [1] |
| YZy81 | *bat1Δ*, isobutanol biosensor (cassette from pYZ16) | CEN.PK2-1C, *his3*::*HIS3*-$P_{LEU1}$-yEGFP-PEST-$T_{ADH1}$-$P_{TPI1}$-$LEU4^{1\text{-}410}$-$T_{PGK1}$, *bat1Δ*::hphMX | This study |
| YZy90 | *pdc1Δ, pdc5Δ, pdc6Δ*, *gal80Δ*, pJLA121-$PDC1^{0202}$ | CEN. PK2-1C, *pdc1Δ, pdc5Δ, pdc6Δ, gal80Δ*::lox71-kanMX-lox66, 2µ *URA3* plasmid ($P_{TFE1}$-*PDC1*-$T_{ACT1}$) | This study |
| YZy91 | *bat1Δ, bat2Δ, ilv6Δ*, isobutanol biosensor (cassette from pYZ16) | CEN.PK2-1C, *his3*::*HIS3*-$P_{LEU1}$-yEGFP-PEST-$T_{ADH1}$-$P_{TPI1}$-$LEU4^{1\text{-}410}$-$T_{PGK1}$, *bat1Δ*::hphMX *bat2Δ*::lox71-kanMX-lox66 *ilv6Δ*::lox71-natMX-lox66 | This study |
| YZy121 | CEN.PK2-1C, isobutanol biosensor (cassette from pYZ16) | CEN.PK2-1C, *his3*::*HIS3*-$P_{LEU1}$-yEGFP-PEST-$T_{ADH1}$-$P_{TPI1}$-$LEU4^{1\text{-}410}$-$T_{PGK1}$ | This study |
| YZy140 | *bat1Δ, leu4Δ, leu9Δ, LEU2* restored | CEN.PK2-1C, *bat1Δ*::hphMX *leu4Δ*::lox71-kanMX-lox66 *leu9Δ*::lox71-natMX-lox66 *leu2::LEU2* | This study |
| YZy148 | *bat1Δ, leu4Δ, leu9Δ, LEU2* restored, modified isopentanol biosensor (cassette from pYZ24) | CEN. PK2-1C, *his3*::*HIS3*-$P_{LEU1}$-yEGFP-$T_{ADH1}$ *bat1Δ*::hphMX *leu4Δ*::lox71-kanMX-lox66 *leu9Δ*::lox71-natMX-lox66 *leu2*::*LEU2* | This study |
| YZy230 | YZy121, cassette from pYZ33 (Strain 1) | YZy121, δ-integration-$P_{TDH3}$-*ILV2*_cHATag-$T_{ADH1}$, $P_{PGK1}$-*ILV3*_cHisTag-$T_{CYC1}$, $P_{TEF1}$-*ILV5*_cMycTag-$T_{ACT1}$ | This study |
| YZy231 | YZy121, cassette from pYZ33 (Strain 2) | YZy121, δ-integration-$P_{TDH3}$-*ILV2*_cHATag-$T_{ADH1}$, $P_{PGK1}$-*ILV3*_cHisTag-$T_{CYC1}$, $P_{TEF1}$-*ILV5*_cMycTag-$T_{ACT1}$ | This study |

| Strain | Description | Genotype (Plasmid contents in parenthesis) | Source |
|---|---|---|---|
| YZy232 | YZy121, cassette from pYZ33 (Strain 3) | YZy121, δ-integration-$P_{TDH3}$-*ILV2*-cHATag-$T_{ADH1}$-$P_{PGK1}$-*ILV3*-cHisTag-$T_{CYC1}$-$P_{TEF1}$-*ILV5*-cMycTag-$T_{ACT1}$ | This study |
| YZy233 | YZy81, cassette from pYZ33 (Strain 1) | YZy81, δ-integration-$P_{TDH3}$-*ILV2*-cHATag-$T_{ADH1}$-$P_{PGK1}$-*ILV3*-cHisTag-$T_{CYC1}$-$P_{TEF1}$-*ILV5*-cMycTag-$T_{ACT1}$ | This study |
| YZy234 | YZy81, cassette from pYZ33 (Strain 2) | YZy81, δ-integration-$P_{TDH3}$-*ILV2*-cHATag-$T_{ADH1}$-$P_{PGK1}$-*ILV3*-cHisTag-$T_{CYC1}$-$P_{TEF1}$-*ILV5*-cMycTag-$T_{ACT1}$ | This study |
| YZy235 | YZy81, cassette from pYZ34 (Strain 1) | YZy81, δ-integration-$P_{TDH3}$-*ILV2*-cHATag-$T_{ADH1}$-$P_{PGK1}$-*ILV3*-cHisTag-$T_{CYC1}$-$P_{TEF1}$-$CoxIV_{MLS}$-*Ll_adhA*$^{RE1}$-cMycTag-$T_{ACT1}$-[$P_{TDH3}$-$CoxIV_{MLS}$-*ARO10*-cHATag-$T_{ADH1}$-$P_{TEF1}$-*ILV5*-cMycTag-$T_{ACT1}$] | This study |
| YZy236 | YZy81, cassette from pYZ34 (Strain 2) | YZy81, δ-integration-$P_{TDH3}$-*ILV2*-cHATag-$T_{ADH1}$-$P_{PGK1}$-*ILV3*-cHisTag-$T_{CYC1}$-$P_{TEF1}$-$CoxIV_{MLS}$-*Ll_adhA*$^{RE1}$-cMycTag-$T_{ACT1}$-[$P_{TDH3}$-$CoxIV_{MLS}$-*ARO10*-cHATag-$T_{ADH1}$-$P_{TEF1}$-*ILV5*-cMycTag-$T_{ACT1}$] | This study |
| YZy311 | YZy91, pYZ127 | YZy91, CEN *URA3* plasmid ($P_{TDH3}$-*ILV6*-$T_{ADH1}$) | This study |
| YZy312 | YZy91, pYZ228 | YZy91, CEN *URA3* plasmid ($P_{TDH3}$-*ILV6*$^{V110E}$-$T_{ADH1}$) | This study |
| YZy313 | YZy363, pYZ127 | YZy363, CEN *URA3* plasmid ($P_{TDH3}$-*ILV6*-$T_{ADH1}$) | This study |
| YZy314 | YZy363, pYZ228 | YZy363, CEN *URA3* plasmid ($P_{TDH3}$-*ILV6*$^{V110E}$-$T_{ADH1}$) | This study |
| YZy363 | YZy81, overexpressing the mitochondrial isobutanol pathway via δ-integration using cassette from pYZ34 | YZy81, δ-integration-$P_{TDH3}$-*ILV2*-cHATag-$T_{ADH1}$-$P_{PGK1}$-*ILV3*-cHisTag-$T_{CYC1}$-$P_{TEF1}$-$CoxIV_{MLS}$-*Ll_adhA*$^{RE1}$-cMycTag-$T_{ACT1}$-[$P_{TDH3}$-$CoxIV_{MLS}$-*ARO10*-cHATag-$T_{ADH1}$-$P_{TEF1}$-*ILV5*-cMycTag-$T_{ACT1}$] | This study |
| YZy418 | *ilv3Δ,* isobutanol biosensor (cassette from pYZ16) | CEN.PK2-1C, *his3*::*HIS3*-$P_{LEU1}$-yEGFP-PEST-$T_{ADH1}$-$P_{TPI1}$-*LEU4*$^{1\text{-}410}$-$T_{PGK1}$, *ilv3Δ*:: lox71-hphMX-lox66 | This study |
| YZy443 | YZy418, *tma29Δ* | YZy418, *tma29Δ*::lox71-kanMX-lox66 | This study |

| Strain | Description | Genotype (Plasmid contents in parenthesis) | Source |
|---|---|---|---|
| YZy447 | YZy443, cytosolic isobutanol pathway (cassette from pYZ196) | YZy443, *ura3Δ*::loxP-*URA3*-loxP-$P_{TDH3}$-*AFT1*-$T_{ADH1}$-[$P_{TEF1}$-*Bs_alsS*-$T_{ACT1}$-$P_{TDH3}$-*Ec_ilvC*$^{P2D1-A1}$-$T_{ADH1}$]-$P_{GAL10}$-*Ll_ilvD*-$T_{ACT1}$ | This study |
| YZy449 | YZy447, *ura3Δ* marker restored | YZy447, *ura3*Δ | This study |
| YZy452 | YZy449, (cassette from pYZ206) | YZy449, δ-integration-$P_{TDH3}$-*Ec_ilvC*$^{P2D1-A1}$-$T_{ADH1}$-[$P_{TEF1}$- *Ec_ilvC*$^{P2D1-A1}$-$T_{ACT1}$] | This study |
| YZy453 | YZy452, pYZ125 | YZy452, empty CEN *URA3* plasmid | This study |
| YZy454 | YZy452, pYZ126 | YZy452, CEN *URA3* plasmid ($P_{TDH3}$-*Ll_ilvD*-$T_{ADH1}$) | This study |
| YZy468 | YZy452, pYZ341 | YZy452, 2μ *URA3* plasmid ($P_{TDH3}$-*Ll_ilvD*-$T_{ADH1}$) | This study |
| YZy469 | YZy452, pYZ353 | YZy452, CEN *URA3* plasmid ($P_{TDH3}$-*Ll_ilvD*$^{I433V}$-$T_{ADH1}$) | This study |
| YZy470 | YZy452, pYZ342 | YZy452, 2μ *URA3* plasmid ($P_{TDH3}$-*Ll_ilvD*$^{I433V}$-$T_{ADH1}$) | This study |
| YZy480 | YZy90, OptoINVRT7, 2μ plasmid pJLA121- *PDC1*$^{0202}$ | YZy90, *his3*::*HIS3*-$P_{TEF1}$-EL222-$T_{CYC1}$-$P_{C120}$-*GAL80*-ODC$^{mut}$-$T_{ACT1}$-[$P_{C120}$-*GAL80*-ODC$^{mut}$-$T_{ACT1}$]-$P_{PGK1}$-*GAL4*-PSD$^{V19L}$-$T_{ADH1}$, 2μ *URA3* plasmid ($P_{TEF1}$-*PDC1*-$T_{ACT1}$) | This study |
| YZy481 | YZy90, OptoINVRT7, 2μ plasmid pJLA121- *PDC1*$^{0202}$ removed | YZy90, *his3*::*HIS3*-$P_{TEF1}$-EL222-$T_{CYC1}$-$P_{C120}$-*GAL80*-ODC$^{mut}$-$T_{ACT1}$-[$P_{C120}$-*GAL80*-ODC$^{mut}$-$T_{ACT1}$]-$P_{PGK1}$-*GAL4*-PSD$^{V19L}$-$T_{ADH1}$ | This study |
| YZy487 | YZy481, isobutanol biosensor integrated to *GAL80* locus (using the cassette from pYZ414) | YZy481, *gal80*::Lox71-natMX6-Lox66-$P_{LEU1}$-yEGFP-PEST-$T_{ADH1}$-$P_{TPI1}$-*LEU4*$^{1-410}$-$T_{PGK1}$ | This study |
| YZy502 | YZy487, OptoEXP *PDC1* and OptoINVRT7 cytosolic isobutanol pathway | YZy487, δ-integration-$P_{C120}$-*PDC1*-$T_{ACT1}$-$P_{TDH3}$-*Ec_ilvC*$^{P2D1-A1}$-$T_{CYC1}$-$P_{TEF1}$-*Ll_ilvD*$^{I433V}$-$T_{TPS1}$-$P_{GAL1-S}$-*Bs_alsS*-$T_{ACT1}$ | This study |

| Strain | Description | Genotype (Plasmid contents in parenthesis) | Source |
|---|---|---|---|
| YZy505 | YZy502, partial cytosolic isobutanol pathway lacking *Bs_alsS* (pYZ350) | YZy502, 2μ *URA3* plasmid ($P_{TDH3}$-*Ec_ilvC*$^{P2D1-A1}$-$T_{CYC1}$-[$P_{TEF1}$-*Ec_ilvC*$^{P2D1-A1}$-$T_{ACT1}$-$P_{TDH3}$-*Ll_ilvD*$^{I433V}$-$T_{ADH1}$]-$P_{PGK1}$-*ARO10*-cHATag-$T_{CYC1}$-$P_{TEF1}$-*Ll_adhA*$^{RE1}$-cMycTag-$T_{ACT1}$) | This study |
| SHy1 | *bat1Δ* | CEN.PK2-1C, *bat1Δ*::hphMX | [2] |
| SHy134 | *bat1Δ, leu4Δ, leu9Δ, LEU2* restored, isopentanol biosensor (cassette from pYZ25) | CEN.PK2-1C, *his3*::*HIS3*-$P_{LEU1}$-yEGFP-$T_{ADH1}$-$P_{TPI1}$-*LEU4*$^{\Delta S547}$-$T_{PGK1}$ *bat1Δ*::hphMX *leu4Δ*::lox71-kanMX-lox66 *leu9Δ*::lox71-natMX-lox66 *leu2::LEU2* | This study |
| SHy158 | SHy134, pYZ125 | SHy134, empty CEN *URA3* plasmid | This study |
| SHy159 | SHy134, JLAb691 | SHy134, CEN *URA3* plasmid ($P_{TDH3}$-*ILV2*_cHATag-$T_{ADH1}$, $P_{PGK1}$-*ILV3*_cHisTag-$T_{CYC1}$, $P_{TEF1}$-*ILV5*_cMycTag-$T_{ACT1}$) | This study |
| SHy176 | SHy134, JLAb705 | SHy134, CEN *URA3* plasmid ($P_{TDH3}$-*ILV1*_cHATag-$T_{ADH1}$) | This study |
| SHy181 | *leu4Δ, leu9Δ, LEU2* restored, *BAT1* restored, isopentanol biosensor (cassette from pYZ25) | CEN. PK2-1C, *his3*::*HIS3*-$P_{LEU1}$-yEGFP-$T_{ADH1}$-$P_{TPI1}$-*LEU4*$^{\Delta S547}$-$T_{PGK1}$ *leu4Δ*::lox71-kanMX-lox66 *leu9Δ*::lox71-natMX-lox66 *leu2*::*LEU2* *trp1*::*TRP1*-$P_{BAT1}$-*BAT1*-$T_{BAT1}$ | This study |
| SHy187 | SHy181, pYZ125 | SHy181, empty CEN *URA3* plasmid | This study |
| SHy188 | SHy181, JLAb691 | SHy181, CEN *URA3* plasmid ($P_{TDH3}$-*ILV2*_cHATag-$T_{ADH1}$, $P_{PGK1}$-*ILV3*_cHisTag-$T_{CYC1}$, $P_{TEF1}$-*ILV5*_cMycTag-$T_{ACT1}$) | This study |
| SHy192 | SHy181, JLAb705 | SHy181, CEN *URA3* plasmid ($P_{TDH3}$-*ILV1*_cHATag-$T_{ADH1}$) | This study |

**Supplementary Table 2. Plasmids used in this study.**

| Plasmid | Description [Brackets indicate inverted orientation] | Source |
|---|---|---|
| pRS416 | Amp$^R$, CEN, URA3 | [3] |
| pRS426 | Amp$^R$, 2μ, URA3 | [4] |
| pYZ1 | Amp$^R$, 2μ, TRP1, $P_{TPI1}$-*LEU4*$^{\Delta S547}$_FLAG-$T_{PGK1}$ | This study |
| pYZ2 | Amp$^R$, 2μ, TRP1, $P_{TPI1}$-*LEU4*$^{1\text{-}410}$_FLAG-$T_{PGK1}$ | This study |
| pYZ12B | Amp$^R$, *HIS3* locus integration vector (His3INT) | [5] |
| pYZ13 | Amp$^R$, 2μ, URA3, $P_{LEU1}$-yEGFP_PEST-$T_{ADH1}$ | This study |
| pYZ14 | Amp$^R$, His3INT, $P_{LEU1}$-yEGFP_PEST-$T_{ADH1}$ | This study |
| pYZ16 | (Isobutanol biosensor)<br>Amp$^R$, His3INT, $P_{LEU1}$-yEGFP_PEST-$T_{ADH1}$-$P_{TPI1}$-*LEU4*$^{1\text{-}410}$_FLAG-$T_{PGK1}$ | This study |
| pYZ17 | Amp$^R$, Lox71-kanMX-Lox66 gene-disruption cassette | [6] |
| pYZ23 | Amp$^R$, δ-integration vector, Lox71-bleMX6-Lox66 | [5] |
| pYZ24 | (Modified isobutanol biosensor without leucine-insensitive Leu4p mutant)<br>Amp$^R$, His3INT, $P_{LEU1}$-yEGFP-$T_{ADH1}$ | This study |
| pYZ25 | (Isopentanol biosensor)<br>Amp$^R$, His3INT, $P_{LEU1}$-yEGFP-$T_{ADH1}$-$P_{TPI1}$-*LEU4*$^{\Delta S547}$_FLAG-$T_{PGK1}$ | This study |
| pYZ33 | (δ-integration-*ILV*s)<br>Amp$^R$, δ-integration-Lox71-ShBle-Lox66-$P_{TDH3}$-*ILV2*-cHATag-$T_{ADH1}$-$P_{PGK1}$-*ILV3*-cHisTag-$T_{CYC1}$-$P_{TEF1}$-*ILV5*-cMycTag-$T_{ACT1}$ | This study |

| | | |
|---|---|---|
| pYZ34 | (δ-integration-ILVs, CoxIV$_{MLS}$-*ARO10*, CoxIV$_{MLS}$-*Ll_adhA*$^{RE1}$)<br>Amp$^{R}$, δ-integration-Lox71-ShBle-Lox66-P$_{TDH3}$-*ILV2*-cHATag-T$_{ADH1}$-P$_{PGK1}$-*ILV3*-cHisTag-T$_{CYC1}$-P$_{TEF1}$-CoxIV$_{MLS}$-*Ll_adhA*$^{RE1}$-cMycTag-T$_{ACT1}$-[P$_{TDH3}$-CoxIV$_{MLS}$-*ARO10*-cHATag-T$_{ADH1}$-P$_{TEF1}$-*ILV5*-cMycTag-T$_{ACT1}$] | [6] |
| pYZ55 | Amp$^{R}$, Lox71-hphMX-Lox66 gene-disruption cassette | [6] |
| pYZ84 | Amp$^{R}$, Lox71-natMX-Lox66 gene-disruption cassette | [6] |
| pYZ113 | (δ-integration-CoxIV$_{MLS}$-*ARO10*, CoxIV$_{MLS}$-*Ll_adhA*$^{RE1}$)<br>Amp$^{R}$, δ-integration-Lox71-ShBle-Lox66-P$_{TDH3}$-CoxIV$_{MLS}$-*ARO10*-cHATag-T$_{ADH1}$-[P$_{TEF1}$-CoxIV$_{MLS}$-*Ll_adhA*$^{RE1}$-cMycTag-T$_{ACT1}$] | This study |
| pYZ125 | Amp$^{R}$, CEN, URA3, P$_{TDH3}$-MCS-T$_{ADH1}$ | This study |
| pYZ126 | Amp$^{R}$, CEN, URA3, P$_{TDH3}$-*Ll_ilvD*-T$_{ADH1}$ | This study |
| pYZ127 | Amp$^{R}$, CEN, URA3, P$_{TDH3}$-*ILV6*-T$_{ADH1}$ | This study |
| pYZ148 | Amp$^{R}$, CEN, URA3, P$_{TDH3}$-*ILV6*$^{V90D/L91F}$-T$_{ADH1}$ | This study |
| pYZ149 | Amp$^{R}$, CEN, URA3, P$_{TDH3}$-*LEU4*-T$_{ADH1}$ | This study |
| pYZ154 | Amp$^{R}$, CEN, URA3, P$_{TDH3}$-*LEU4*$^{\Delta S547}$-T$_{ADH1}$ | This study |
| pYZ155 | Amp$^{R}$, CEN, URA3, P$_{TDH3}$-*LEU4*$^{1\text{-}410}$-T$_{ADH1}$ | This study |
| pYZ196 | (Cytosolic isobutanol pathway containing P$_{GAL10}$-*Ll_ilvD*)<br>Amp$^{R}$, Ura3 Locus integration-LoxP-*URA3*-LoxP- P$_{TDH3}$-*AFT1*-T$_{ADH1}$-[P$_{TEF1}$-*Bs_alsS*-T$_{ACT1}$-P$_{TDH3}$-*Ec_ilvC*$^{P2D1\text{-}A1}$-T$_{ADH1}$]-P$_{GAL10}$-*Ll_ilvD*-T$_{ACT1}$ | This study |
| pYZ206 | (δ-integration-two copies of *Ec_ilvC*$^{P2D1\text{-}A1}$)<br>Amp$^{R}$, δ-integration-Lox71-ShBle-Lox66-P$_{TDH3}$-*Ec_ilvC*$^{P2D1\text{-}A1}$-T$_{ADH1}$-[P$_{TEF1}$-*Ec_ilvC*$^{P2D1\text{-}A1}$-T$_{ACT1}$] | This study |
| pYZ223 | Amp$^{R}$, Lox71-natMX6-Lox66, *GAL80* Locus Integration vector (Gal80INT-Lox71-natMX6-Lox66) | This study |

| | | |
|---|---|---|
| pYZ228 | Amp$^R$, CEN, URA3, P$_{TDH3}$-*ILV6*$^{V110E}$-T$_{ADH1}$ | This study |
| pYZ341 | Amp$^R$, 2μ, URA3, P$_{TDH3}$-*Ll_ilvD*-T$_{ADH1}$ | This study |
| pYZ342 | Amp$^R$, 2μ, URA3, P$_{TDH3}$-*Ll_ilvD*$^{I433V}$-T$_{ADH1}$ | This study |
| pYZ350 | (Partial cytosolic isobutanol pathway containing *ARO10, Ll_adhA*$^{RE1}$, *Ll_ilvD*$^{I433V}$, and extra copies of *Ec_ilvC*$^{P2D1\text{-}A1}$; lacking *Bs_alsS*)<br>Amp$^R$, 2μ, URA3, P$_{TDH3}$-*Ec_ilvC*$^{P2D1\text{-}A1}$-T$_{CYC1}$-[P$_{TEF1}$-*Ec_ilvC*$^{P2D1\text{-}A1}$-T$_{ACT1}$]-[P$_{TDH3}$-*Ll_ilvD*$^{I433V}$-T$_{ADH1}$]-P$_{PGK1}$-*ARO10*-cHATag-T$_{CYC1}$-P$_{TEF1}$-*Ll_adhA*$^{RE1}$-cMycTag-T$_{ACT1}$ | This study |
| pYZ353 | Amp$^R$, CEN, URA3, P$_{TDH3}$-*Ll_ilvD* $^{I433V}$-T$_{ADH1}$ | This study |
| pYZ383 | Amp$^R$, 2μ, URA3, P$_{TEF1}$-*Ll_ilvD*$^{I433V}$-T$_{TPS1}$ | This study |
| pYZ384 | Amp$^R$, 2μ, URA3, P$_{Gal1\text{-}S}$-*Bs_alsS*-T$_{ACT1}$ | This study |
| pYZ414 | Amp$^R$, Gal80INT-Lox71-natMX6-Lox66-Isobutanol biosensor (P$_{LEU1}$-yEGFP_PEST-T$_{ADH1}$-P$_{TPI1}$-*LEU4*$^{1\text{-}410}$_FLAG-T$_{PGK1}$) | This study |
| pYZ417 | (δ-integration-OptoEXP-*PDC1*-OptoINVRT7-cytosolic isobutanol pathway)<br>Amp$^R$, δ-integration-Lox71-ShBle-Lox66-OptoEXP-*PDC1* (P$_{C120}$-*PDC1*-T$_{ACT1}$)-OptoINVRT7-cytosolic isobutanol pathway (P$_{TDH3}$- *Ec_ilvC*$^{P2D1\text{-}A1}$-T$_{CYC1}$-P$_{TEF1}$-*Ll_ilvD*$^{I433V}$-T$_{TPS1}$-P$_{Gal1\text{-}S}$-*Bs_alsS*-T$_{ACT1}$) | This study |
| EZ-L235 | (δ-integration-OptoEXP-*PDC1*)<br>Amp$^R$, δ-integration-Lox71-ShBle-Lox66-OptoEXP-*PDC1* (P$_{C120}$-*PDC1*-T$_{ACT1}$) | [5] |
| EZ-L439 | (OptoINVRT7)<br>Amp$^R$, His3INT-P$_{TEF1}$-EL222-T$_{CYC1}$-P$_{C120}$-*GAL80*-ODC$^{mut}$-T$_{ACT1}$-[P$_{C120}$-*GAL80*-ODC$^{mut}$-T$_{ACT1}$]-P$_{PGK1}$-*GAL4*-PSD$^{V19L}$ -T$_{ADH1}$ | [7] |
| pAG26 | Amp$^R$, Plasmid containing hphMX gene-disruption cassette | [8] |
| pJA123 | Amp$^R$, 2μ, URA3, P$_{TDH3}$-*ILV2*-cHATag-T$_{ADH1}$-P$_{PGK1}$-*ILV3*-cHisTag-T$_{CYC1}$-P$_{TEF1}$-*ILV5*-cMycTag-T$_{ACT1}$ | [9] |

| | | |
|---|---|---|
| pJA182 | $Amp^R$, 2µ, URA3, $P_{TDH3}$-*ILV2*-HA-$T_{ADH1}$-$P_{PGK1}$-*ILV3*-cHisTag-$T_{CYC1}$-$P_{TEF1}$-$CoxIV_{MLS}$-*Ll_adhA*$^{RE1}$-cMycTag-$T_{ACT1}$-[$P_{TDH3}$-$CoxIV_{MLS}$-*ARO10*-cHATag-$T_{ADH1}$-$P_{TEF1}$-*ILV5*-cMycTag-$T_{ACT1}$] | [9] |
| pJA248 | $Amp^R$, Plasmid containing yEGFP and PEST protein degradation tag | This study |
| pJLA121-*PDC1*$^{0202}$ | $Amp^R$, 2µ, URA3, $P_{TFE1}$-*PDC1*-$T_{ACT1}$ | [5] |
| JLAb23 | $Amp^R$, 2µ, URA3, pRS426-$P_{TPI1}$-MCS_FLAG-$T_{PGK1}$ | This study |
| JLAb131 | $Amp^R$, 2µ, URA3, pJLA121$^{0103}$- $P_{TDH3}$-MCS-$T_{ADH1}$ | This study |
| JLAb581 | $Amp^R$, 2µ, URA3, $P_{TDH3}$-$CoxIV_{MLS}$-*ARO10*-cHATag-$T_{ADH1}$-[$P_{TEF1}$- $CoxIV_{MLS}$-*Ll_adhA*$^{RE1}$-cMycTag-$T_{ACT1}$] | This study |
| JLAb691 | $Amp^R$, CEN, URA3, $P_{TDH3}$-*ILV2*_cHATag-$T_{ADH1}$-$P_{PGK1}$-*ILV3*_cHisTag-$T_{CYC1}$-$P_{TEF1}$-*ILV5*_cMycTag-$T_{ACT1}$ | This study |
| JLAb705 | $Amp^R$, CEN, URA3, $P_{TDH3}$-*ILV1*_cHATag-$T_{ADH1}$ | This study |
| pSH63 | $Amp^R$, CEN, TRP1, $P_{GAL1}$_Cre_$T_{CYC1}$ | [10] |

**Supplementary Table 3.** Amino acid mutations found in the sequenced Ilv6p variants.

| Mutants | Mutations |
|---|---|
| **Derived from wild-type *ILV6* random mutagenesis library** | |
| ILV6_mutant_1 | N104S |
| ILV6_mutant_2 | S71P_N86T |
| ILV6_mutant_3 | N86T |
| ILV6_mutant_4 | V90A |
| ILV6_mutant_5 | M21I_Y34C_N223Y |
| ILV6_mutant_6 | V110E |
| ILV6_mutant_7 | N86Y |
| ILV6_mutant_8 | L91S |
| ILV6_mutant_9 | N104S_E133V_S184F_G233E |
| ILV6_mutant_10 | M21I_N57S_V79D_E133G_L161Q_T185A_N186S_D224Y_K284E_E288K |
| **Derived from *ILV6*$^{V90D/L91F}$ random mutagenesis library** | |
| ILV6_mutant_11 | V90D_L91F_D292N |
| ILV6_mutant_12 | V90D_L91F_H219L |
| ILV6_mutant_13 | V90D_L91F_N112S |
| ILV6_mutant_14 | V90D_L91F_T185A |
| ILV6_mutant_15 | N86D_V90D_L91F |
| ILV6_mutant_16 | N86S_V90D_L91F_K202M |
| ILV6_mutant_17 | V90D_L91F_F174S |
| ILV6_mutant_18 | V90D_L91F_N153Y_E198G |
| ILV6_mutant_19 | P47T_Q85R_V90D_L91F_H180R |
| ILV6_mutant_20 | V90D_L91F_V267A |
| ILV6_mutant_21 | V90D_L91F_Q126L_S286R |
| ILV6_mutant_22 | V90D_L91F_H180Y_I239V |
| ILV6_mutant_23 | R11H_V84M_V90D_L91F_F174I |
| ILV6_mutant_24 | V90D_L91F_Q193R |

**Supplementary Table 4.** Amino acid mutations found in the sequenced Leu4p variants.

| Mutants | Mutations |
|---|---|
| **Derived from mutagenesis library of full-length *LEU4*** | |
| LEU4_mutant_1 | Y203F_V425D_N515I |
| LEU4_mutant_2 | K25N_A60T_T316N_L330M_D433G_K467R_F497L_D509G_N515D_L529M_A552V_I602L |
| LEU4_mutant_3 | N72S_K97R_S126G_R344H_R495G_N515D |
| LEU4_mutant_4 | V446A_G544C_N593Y |
| LEU4_mutant_5 | H541R |
| LEU4_mutant_6 | E86D_K191N_K374R_A445T_S481R_N515I_A568V_S601A |
| LEU4_mutant_7 | T287A_P400L_N515H_N537T |
| LEU4_mutant_8 | A144T_N240S_D578N |
| LEU4_mutant_9 | K90M_Q439R_S542P |
| LEU4_mutant_10 | R428G_N486D |
| LEU4_mutant_11 | K51R_E233K_F377I_L427S_K458E_F497I_E577D |
| LEU4_mutant_12 | R392G_S459L_V584A |
| LEU4_mutant_13 | K51R_A182V_V198T_A551V |
| **Derived from mutagenesis library of regulatory domain** | |
| LEU4_mutant_14 | Y485N |
| LEU4_mutant_15 | A450S_D451G_D578E |
| LEU4_mutant_16 | T590I_P603S |
| LEU4_mutant_17 | K489E_V573A |
| LEU4_mutant_18 | Y538N |
| LEU4_mutant_19 | Q447R_Q478H_G516D |
| LEU4_mutant_20 | V584E |
| LEU4_mutant_21 | Q439H_D581G |
| LEU4_mutant_22 | R436K_S443Y |
| LEU4_mutant_23 | D564E_T590I |
| LEU4_mutant_24 | T590I |

**Supplementary Table 5.** Occurrence and description of the most relevant mutations found in the Leu4p variants.

| Position | Number of occurrences | Variants | Single or multiple mutant variant# | Number of substitutions | Type of substitutions |
|---|---|---|---|---|---|
| **N515** | 5 | 1, 2, 3, 6, 7 | Multiple | 3 | I, D, H |
| **T590** | 3 | 16, 23, 24 | Single and Multiple | 1 | I |
| **K51** | 2 | 11, 13 | Multiple | 1 | R |
| **Q439** | 2 | 9, 21 | Multiple | 2 | R, H |
| **F497** | 2 | 2, 11 | Multiple | 2 | L, I |
| **D578** | 2 | 8, 15 | Multiple | 2 | N, E |
| **V584** | 2 | 12, 20 | Single and Multiple | 2 | A, E |
| **Y485** | 1 | 14 | Single | 1 | N |
| **Y538** | 1 | 18 | Single | 1 | N |
| **H541** | 1 | 5 | Single | 1 | R |

### Mutations observed in variants with only one mutation (Single), as one of the mutations in variants with multiple mutations (Multiple), or in both types of variants (Single and Multiple).

**Supplementary Table 6.** Published kinetic parameters of *Bs*_AlsS, *Ec*_IlvC$^{P2D1-A1}$, and *Ll*_IlvD.

| Enzymes | Specific activity (U/mg) | $K_m$ (mM) | $k_{cat}$ ($s^{-1}$) | $k_{cat}/K_m$ | Assay conditions | References |
|---|---|---|---|---|---|---|
| *Bs*_AlsS | 8.28 | 13.6 ± 0.8 | 121 ± 13 | 8.9 + 1.1 | pH 7.0, 37°C | [11] |
| *Ec*_IlvC$^{P2D1-A1}$ | n.a. | n.a. | 4.3 ± 0.3 | | pH 7.0, n.a. | [12] |
| *Ll*_IlvD | 0.62 ±0.01* | n.a. | n.a. | | n.a. | [13] |

* The highest value reported. n.a.: not available

**Supplementary Table 7.** Amino acid mutations found in the sequenced *Ll*_IlvD variants.

| **Mutants** | **Mutations** |
|---|---|
| *Ll*_ilvD_mutant_1 | I433V |
| *Ll*_ilvD_mutant_2 | V12A, S189P, H439R |
| *Ll*_ilvD_mutant_3 | K535R |
| *Ll*_ilvD_mutant_4 | K16R |
| *Ll*_ilvD_mutant_5 | E13G, K345M, I514N |
| *Ll*_ilvD_mutant_6 | I154V, I312T |

**Supplementary Table 8.** Oligonucleotides used in this study.

| Oligo Name | Sequence | Description |
|---|---|---|
| Yfz_Oli31 | GCTGGAGCTCACCGGTATACCCGGAATATGAACCACAGTACATCATATTAAGACGTAGT | $P_{LEU1}$_Gibson_primer_F |
| Yfz_Oli32 | GAATAATTCTTCACCTTTAGACATGATTTAAAACAGCAAATAATAAAAATCGATAGCGAC | $P_{LEU1}$_Gibson_primer_R |
| Yfz_Oli33 | CGATTTTTATTATTTGCTGTTTTAAATCATGTCTAAAGGTGAAGAATTATTCACTGGTG | yEGFP_PEST-Gibson_Primer_F |
| Yfz_Oli34 | TCGCTGATCATTACTCGAGGTCGACCTATATTACTTGGGTATTGCCCATACC | yEGFP_PEST-Gibson_Primer_R |
| Yfz_Oli35 | CATGGCTAGCGTTAAAGAGAGTATTATTGC | NheI-ScLEU4$^{1-410}$ Primer-F |
| Yfz_Oli36 | ATAATCCTCGAGGACAGCTTCGTAATCACGGC | XhoI-ScLEU4$^{1-410}$ Primer-R |
| Yfz_Oli37 | CTGAGCGGCCGCTAAAATCATGGCTAGCGTTAAAGAGAGTATTATTGC | NotI-KOZAK-NheI-ScLEU4$^{WT}$ Primer-F |
| Yfz_Oli38 | ATAATCCTCGAGTGCAGAGCCAGATGCCGCAGCATTCTTA | XhoI-ScLEU4$^{WT}$ Primer-R |
| Yfz_Oli59 | TCGACACGCGTTTATTT | Annealed oligo cloning linker SalI_MluI_BsrGI_F |
| Yfz_Oli60 | GTACAAATAAACGCGTG | Annealed oligo cloning linker SalI_MluI_BsrGI_R |
| Yfz_Oli198 | CCGCTAAAATCATGGCTAGC | Error-prone PCR universal_Primer_F for full ORF subcloned into pYZ125 |
| Yfz_Oli242 | CATAAATCATAAGAAATTCGCTGATCATTACTCGAG | Error-prone PCR universal_Primer_R for full ORF subcloned into pYZ12s |
| Yfz_Oli243 | GCCGCTTGGGTTATTTTGAGATCT | Error-prone PCR Primer_F for Leu-regulatory domain (Leu430-Ala618, BglII_ScLEU4_XhoI) of ScLEU4 |

| | | |
|---|---|---|
| Yfz_Oli345 | ATCATGGCTAGCGAATTTAAGTACAACG GTAAGGTC | NheI_*Ll_ilvD*_F |
| Yfz_Oli346 | ATACATCTCGAGTCATTACAAGTCGGTAA CACAACCTTCAG | XhoI_*Ll_ilvD*_R |
| Jla_oli234 | GATCGCTAGCCTGAGATCGTTATTGCAAA GC | NheI_ScILV6_F |
| Jla_oli235 | AATTCTCGAGACCAGGTGGTAGTTGGGA AATG | XhoI_ScILV6 _R |
| Jla_oli276 | GAATCGCTAGCGTTAAAGAGAGTATTAT TGCTCTTGCTGAGC | NheI_ScLEU4_F |
| Jla_oli276R | TCATTACTCGAGTGCAGAGCCAGATGCC GCAGCATTCTTA | XhoI_ScLEU4_R |
| Jla_oli280 | CAGAGCATTCTCTAGGTTCTGGTTCTACG CAAGCTGCTTCTTACATCC | LEU4$^{\Delta S547}$_site-directed mutagenesis Primer-F |
| Jla_oli281 | GGATGTAAGAAGCAGCTTGCGTAGAACC AGAACCTAGAGAATGCTCTG | LEU4$^{\Delta S547}$_site-directed mutagenesis Primer-R |
| Yfz_KO71 | CCAGCGTATACAATCTCGATAGTTGGTTT CCCGTTCTTTCCACTCCCGTCtacgctgcaggtc gacaacc | ScGAL80_KO_F |
| Yfz_KO72 | GTTTTTATAACGTTCGCTGCACTGGGGGC CAAGCACAGGGCAAGATGCTTccactagtggat ctgatatcacc | ScGAL80_KO_R |
| Yfz_KO187 | AGAAAAAAAAGGATTCTCACACTAGAAG TTTACTGTAGACTTTTTCCTTACAAAAAG ACAAGGAACAATCtacgctgcaggtcgacaacc | ScLEU4_KO_F |
| Yfz_KO188 | AGGAAAGGAAGTAAATAAATAAGTATAG AAATAAATAGAAGCGAATAAGTCCTGAA ATACAGAAAAGTTCctagtggatctgatatcacc | ScLEU4_KO_R |
| Yfz_KO189 | ACTACATGTTTTCGTTAGAATAAATCACC CTATAAACGCAAAATCAGCTAGAACCTT AGCATACTAAAACtacgctgcaggtcgacaacc | ScBAT1_KO_F |
| Yfz_KO190 | AACAGATCCTCTGAGAGGAATTCTCGTTT TTTTTTTTTGGGGGGGGAGGGGATGTTTA CCTTCATTATCActagtggatctgatatcacc | ScBAT1_KO_R |
| Yfz_KO229 | TGTTTTCGGCTTATAAGGGTCTTCTCCTT AGGATAATACTATCGGCACATTATCATTT AGCCGCGTAGCCtacgctgcaggtcgacaacc | ScLEU9_KO_F |

| | | |
|---|---|---|
| Yfz_KO230 | TTTTCTGTGCCATTTATAAATAAAAATAC ATATATATATAACATGAGTAATCATAAG CTACTCCTTTCTActagtggatctgatatcacc | ScLEU9_KO_R |
| Yfz_KO266 | ATTGTAGCGCCTGTAATCTTTAGTAACGG ATTCTTGTATTTTTTTGTAAACAGCCAAG AAAAAAGTAGAGtacgctgcaggtcgacaacc | ScILV3_KO_F |
| Yfz_KO267 | TGCGAACAAAAAAGATGATGGAAAAGG AGAATCTCTATATATATATTCATCGATTG GGGCCTATAATGCActagtggatctgatatcacc | ScILV3_KO_R |
| Yfz_KO278 | AAGCTCACTAGTAAAGGCGGGAAATAGA ACATTGAGAACGTATTTTGATAtacgctgcagg tcgacaacc | ScTMA29_KO_F |
| Yfz_KO279 | AAAGTCTTACATGTATAAAAAGTATACA GATTTACTTAGTTTAGCTAGGTctagtggatctg atatcacc | ScTMA29_KO_R |
| Jla_KO1 | TATTTTCTACTCATAACCTCACGCAAAAT AACACAGTCAAATCAATCAAAtacgctgcaggt cgacaacc | ScPDC1_KO_F |
| Jla_KO2 | TACATAAAAATGCTTATAAAACTTTAACT AATAATTAGAGATTAAATCGCccactagtggat ctgatatcacc | ScPDC1_KO_R |
| Jla_KO3 | CATAATCAATCTCAAAGAGAACAACACA ATACAATAACAAGAAGAACAAAtacgctgca ggtcgacaacc | ScPDC5_KO_F |
| Jla_KO4 | AAAGTAAAAAAATACACAAACGTTGAAT CATGAGTTTTATGTTAATTAGCccactagtggat ctgatatcacc | ScPDC5_KO_R |
| Jla_KO5 | AGTATAAATAAAAAACCCACGTAATATA GCAAAAACATATTGCCAACAAAtacgctgcag gtcgacaacc | ScPDC6_KO_F |
| Jla_KO6 | TAAGTTTATTTATTTGCAACAATAATTCG TTTGAGTACACTACTAATGGCccactagtggatc tgatatcacc | ScPDC6_KO_R |
| Jla_KO22 | AAAATTTTAGAAATTTAAGGGAAAGCAT CTCCACGAGTTTTAAGAACGATtacgctgcagg tcgacaacc | ScBAT2_KO |
| Jla_KO23 | AGTTTTATTCTTTTTAACTTTTAATTACTT TACGTAGCAATAGCGATACTccactagtggatct gatatcacc | ScBAT2_KO |

**Supplementary Figures**

**Supplementary Figure 1**

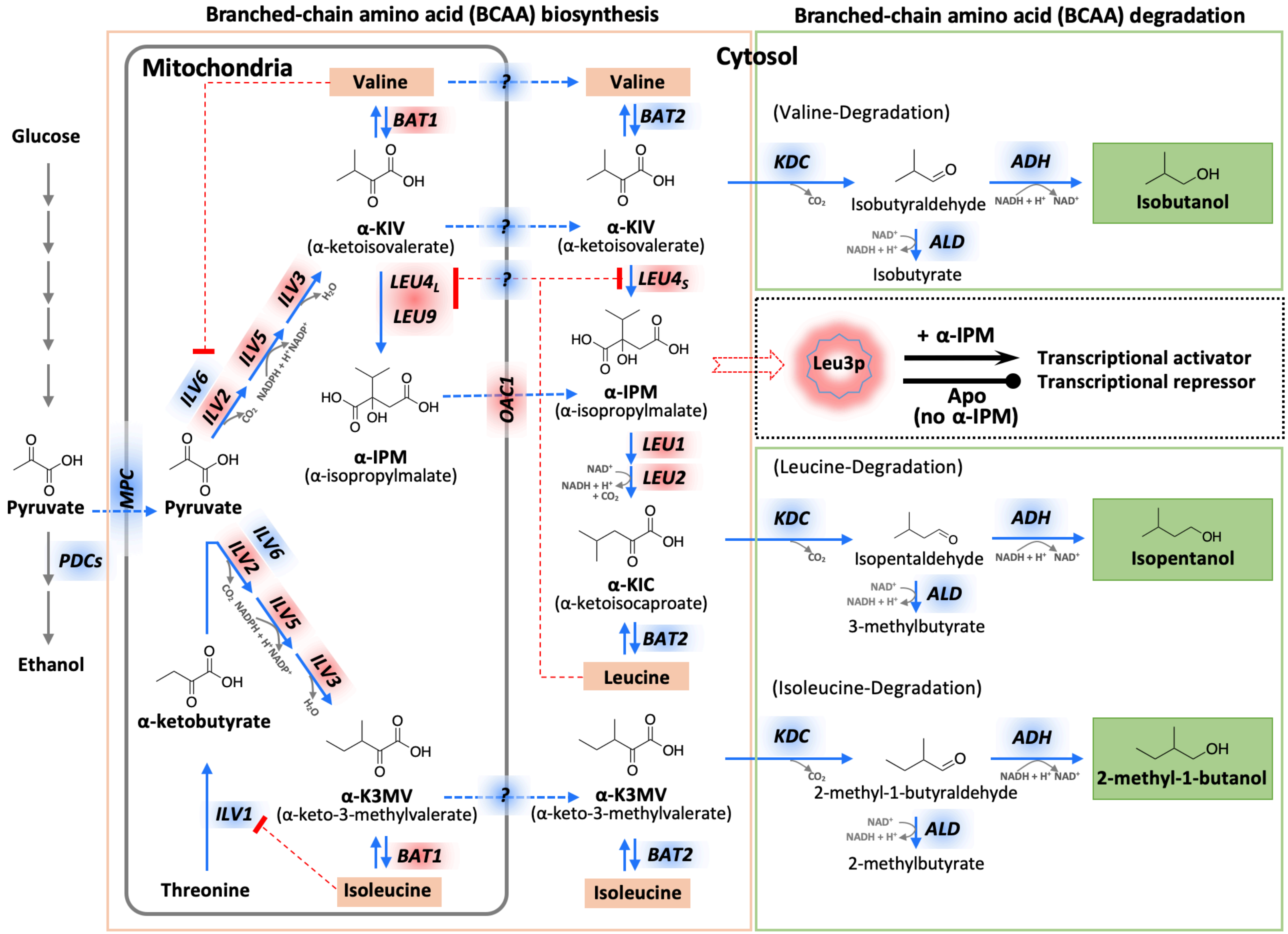

**Supplementary Figure 1. Pathways for branched-chain amino acid (BCAA) and branched-chain higher alcohol (BCHA) biosynthesis in *Saccharomyces cerevisiae*.** Isobutanol, isopentanol, and 2-methyl-1-butanol biosynthesis are derived from the biosynthesis and degradation of valine, leucine, and isoleucine, respectively. The upstream pathway for isobutanol production (valine biosynthesis) consists of three enzymes natively localized in mitochondria (orange rectangle): acetolactate synthase (ALS, encoded by *ILV2*), ketol-acid reductoisomerase (KARI, encoded by *ILV5*), and dehydroxyacid dehydratase (DHAD, encoded by *ILV3*)[14]. Ilv2p, Ilv3p, and Ilv5p convert two molecules of pyruvate to the valine precursor α-ketoisovalerate (α-KIV), which is exported to the cytosol by one or more unknown α-KIV carrier(s). The native localization of the Ehrlich valine degradation occurs in the cytosol, where α-KIV is converted to isobutanol through the Ehrlich BCAA degradation pathway[15] (green rectangle), comprised of α-ketoacid decarboxylases (α-KDCs) and alcohol dehydrogenases (ADHs). The conversion between α-KIV and valine is catalyzed by mitochondrial and cytosolic branched-chain amino acid aminotransferases (encoded by *BAT1* and *BAT2,* respectively). In the upstream pathway for isopentanol production (leucine biosynthesis), α-KIV is converted to α-isopropylmalate (α-IPM) by α-IPM synthases located in mitochondria (encoded by the short $LEU4_S$ and *LEU9*) and the cytosol (encoded by the long $LEU4_L$). Subsequently, α-IPM is converted in the cytosol to β-IPM by isopropylmalate isomerase (encoded by *LEU1*) and then to α-ketoisocaproate (α-KIC) by β-IPM dehydrogenase (encoded by *LEU2*). This α-KIC precursor is then converted to leucine by Bat2p. Alternatively, α-KIC is converted to isopentanol in the cytosol via the BCAA Ehrlich degradation pathway[15]. The upstream pathway for 2-methyl-1-butanol (isoleucine biosynthesis) consists of the same mitochondrial enzymes involved in valine and leucine biosynthesis, Ilv2p, Ilv5p, and Ilv3p; except that for isoleucine biosynthesis, Ilv2p catalyzes the condensation of one pyruvate and one α-ketobutyrate, produced by threonine deaminase (encoded by *ILV1*), instead of two pyruvate molecules as in the biosynthesis of valine and leucine. The subsequent reactions catalyzed by Ilv5p and Ilv3p result in the production of the isoleucine precursor α-keto-3-methylvalerate (α-K3MV), which is transaminated to isoleucine in the mitochondria and the cytosol by Bat1p and Bat2p, respectively. Alternatively, α-K3MV is converted to 2-methyl-1-butanol through the Ehrlich BCAA degradation pathway (green rectangle) in the cytosol. The enzymatic activities of Ilv6p, Leu4p/Leu9p, and Ilv1p are negatively regulated by valine, leucine, and isoleucine, respectively, as indicated with red dashed lines. Leu3p, a dual-function transcriptional regulator, regulates genes (highlighted in red) involved in BCAA biosynthesis. It acts as a transcriptional activator in the presence of α-isopropylmalate (α-IPM) and a repressor in its absence. Genes not known to be regulated by Leu3p are labeled in blue. *PDC*s: pyruvate decarboxylase isozymes; *MPC*: mitochondrial pyruvate carriers; *OAC1*: mitochondrial α-IPM transporter; *ALD*: aldehyde dehydrogenase.

**Supplementary Figure 2**

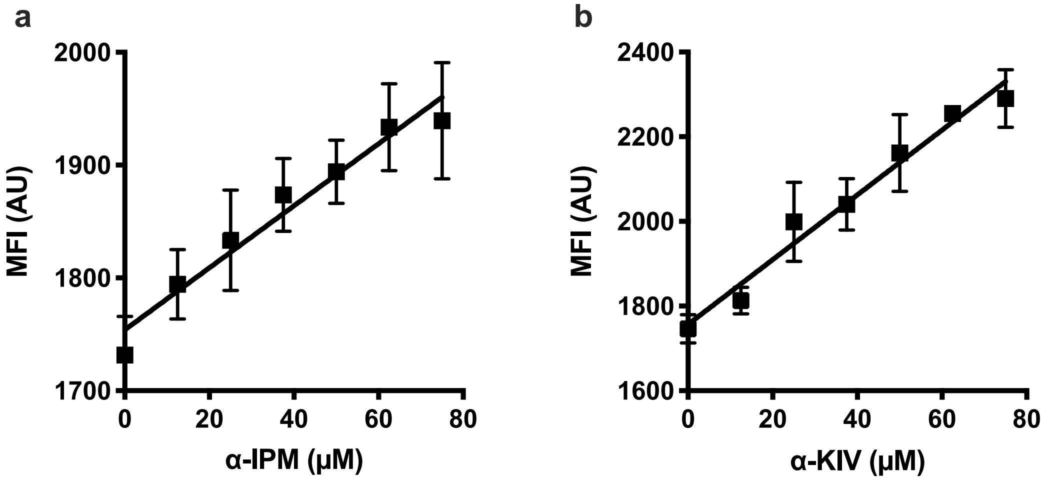

**Supplementary Figure 2. Response of the isobutanol biosensor to α-IPM and α-KIV.** The isobutanol biosensor responds linearly to increasing concentrations of α-isopropylmalate (α-IPM, **a**) and α-ketoisovalerate (α-KIV, **b**) supplemented in the media. MFI: median fluorescence intensity. Error bars represent the standard deviation of at least three biological replicates.

#### Supplementary Figure 3

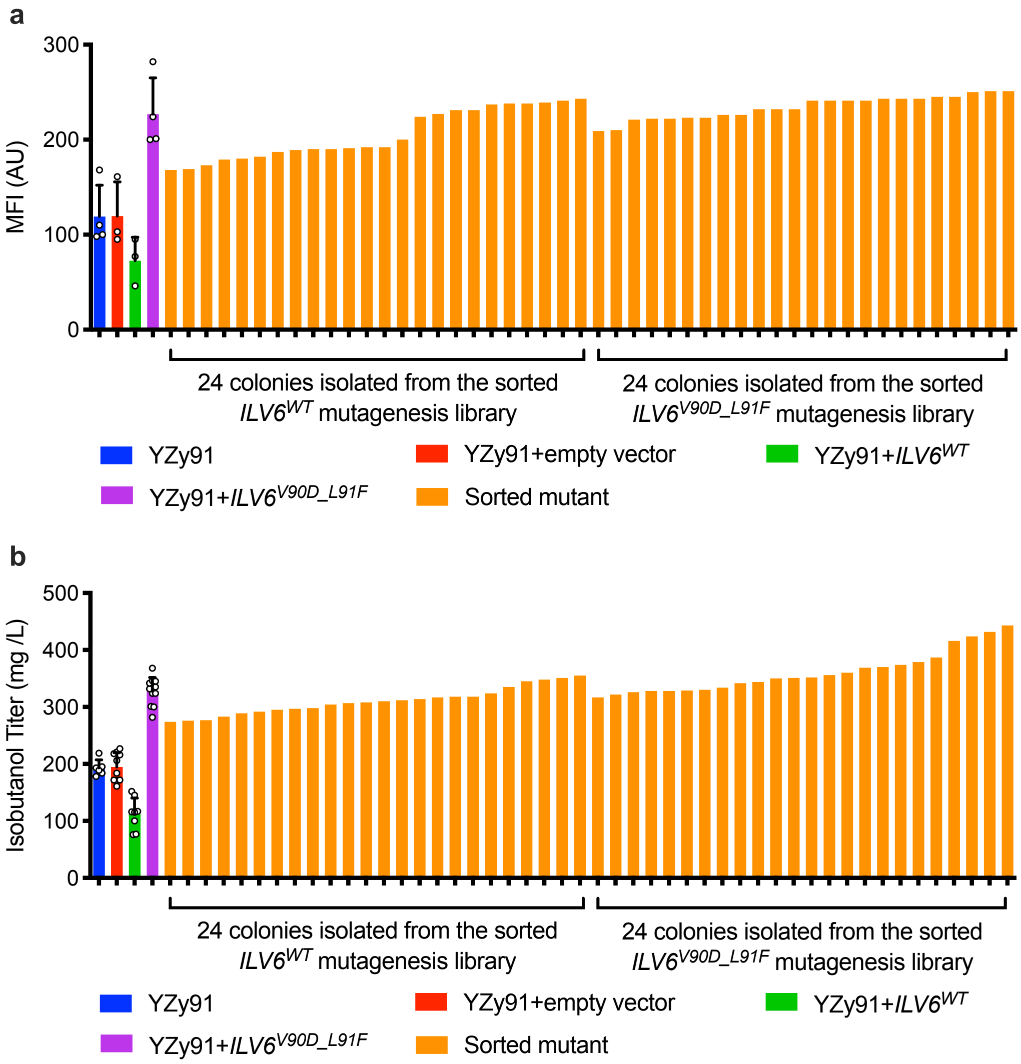

**Supplementary Figure 3. Screen of 48 random colonies sorted from mutagenesis libraries of the wild-type *ILV6* and valine-insensitive *ILV6*$^{V90D\_L91F}$ mutant.** (**a**) Flow cytometry measurements of the GFP median fluorescence intensity (MFI) of 48 sorted strains (orange) measured after 13h of growth in media containing four times more valine (2.4 mM) than the usual synthetic defined medium (24 colonies were isolated from the sorted wild-type *ILV6* mutagenesis library, and 24 colonies were isolated from the sorted *ILV6*$^{V90D\_L91F}$ mutagenesis library). MFI of the basal strain YZy91 (*ilv6Δ bat1Δ bat2Δ*) with (red) or without (blue) an empty vector, or transformed with plasmids containing *ILV6*$^{WT}$ (green) or *ILV6*$^{V90D\_L91F}$ (purple), are shown as controls. (**b**) Isobutanol titers after 48h fermentations with the same 48 sorted strains (orange) and controls (red, blue, purple, and green) shown in (**a**). Error bars of the control strains represent the standard deviation of at least three biological replicates.

**Supplementary Figure 4**

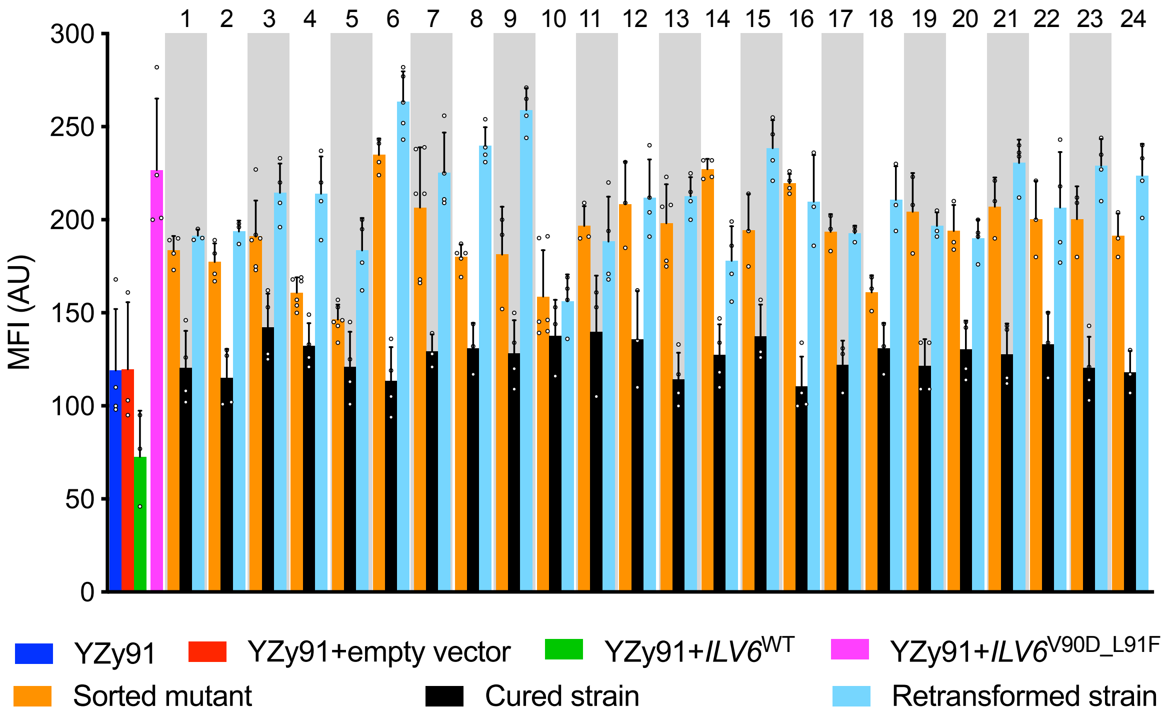

**Supplementary Figure 4. Confirmation that isolated *ILV6* variants with unique sequences enhance GFP fluorescence signal from the isobutanol biosensor.** Flow cytometry measurements of the GFP median fluorescence intensity (MFI) of sorted strains with unique *ILV6* sequences (orange), plasmid-cured derivatives (black), and strains obtained from retransforming YZy91 with each unique plasmid isolated from the corresponding sorted strains (cyan), measured after 13h of growth in media containing four times more valine (2.4 mM) than the usual synthetic defined medium. GFP fluorescence measured with the basal strain YZy91 (*ilv6Δ bat1Δ bat2Δ*) with (red) or without (blue) an empty vector, or transformed with plasmids containing wild-type *ILV6* (*ILV6$^{WT}$*, green), or *ILV6*$^{V90D_L91F}$ (magenta), are shown as controls. The numbers on the upper x-axis identify each of the strains harboring screened mutants with unique *ILV6* sequences. *ILV6* mutants 1 – 10 were isolated from the sorted *ILV6*$^{WT}$ mutagenesis library. *ILV6* mutants 11 – 24 were isolated from the sorted *ILV6*$^{V90D\_L91F}$ mutagenesis library. Error bars represent the standard deviation of three biological replicates.

#### Supplementary Figure 5

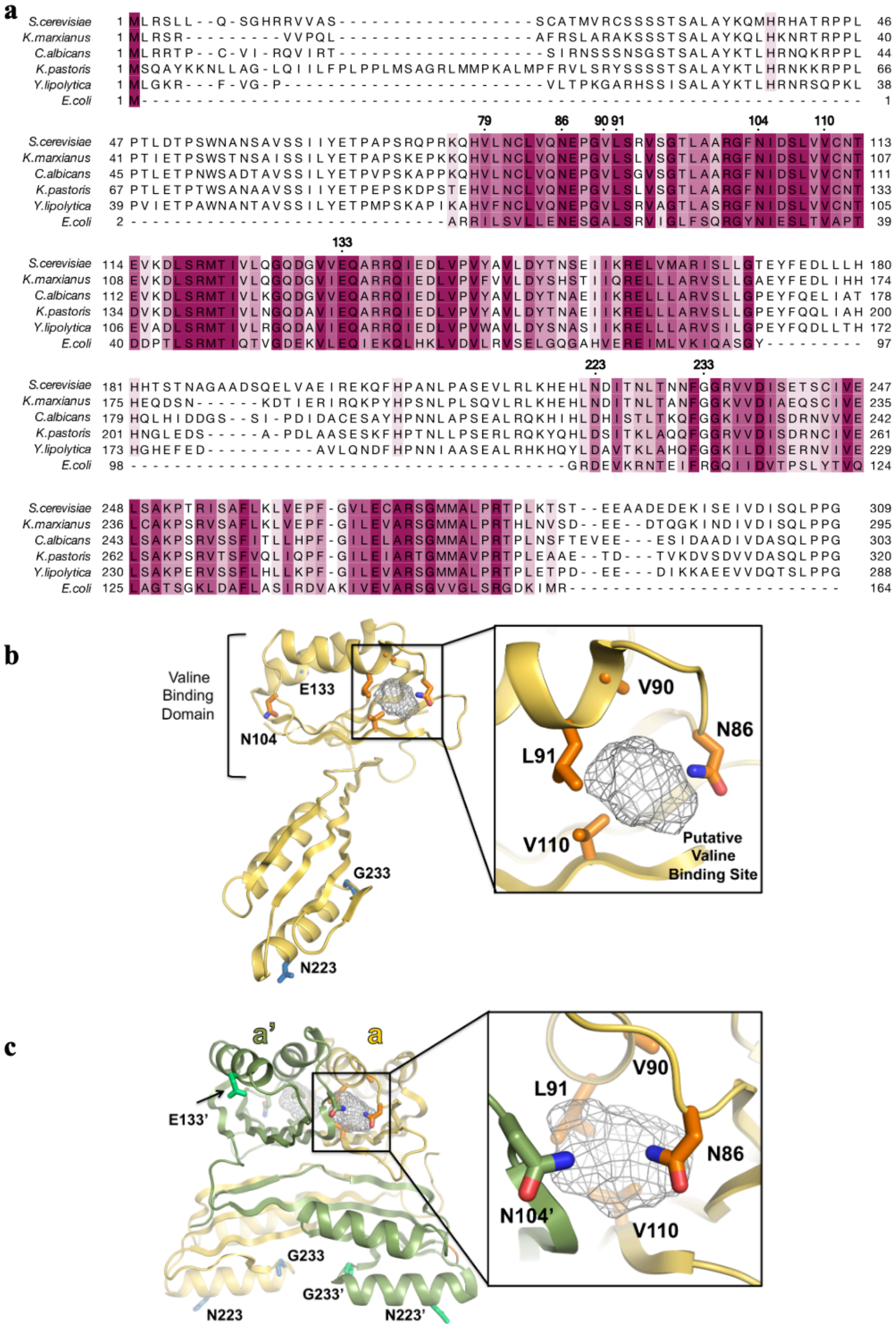

**Supplementary Figure 5. Sequence and structural analysis of yeast Ilv6p and its isolated mutants.** (**a**) Alignment of the amino acid sequence of *E.coli* IlvH (acetohydroxyacid synthase regulatory subunit) with Ilv6p sequences from *Saccharomyces cerevisiae*, *Kluyveromyces marxianus*, *Candida albicans*, *Komagataella pastoris*, and *Yarrowia lipolytica*. The color intensity reflects the level of conservation of each residue. Key residues mutated in *ILV6*, derived from the *ILV6* wild-type mutagenesis library (Supplementary Table 3), that enhance isobutanol production and whose positions are present in *E.coli* IlvH (N86, V90, L91, N104, V110, N223, E133, and G233) are labeled on the top of the alignment (most of them are highly conserved except G233). (**b, c**) Key substituted residues above mapped onto the crystal structure of the regulatory subunit of acetohydroxyacid synthase, IlvH, from *E. coli* (pdb: 2f1f), shown on the structure of the monomer (**b**), and from a different angle on the structure of the dimer (**c**). Close-up views of the putative valine-binding sites (gray mesh) and the key residues that make them are shown in the side-boxes.

**Supplementary Figure 6**

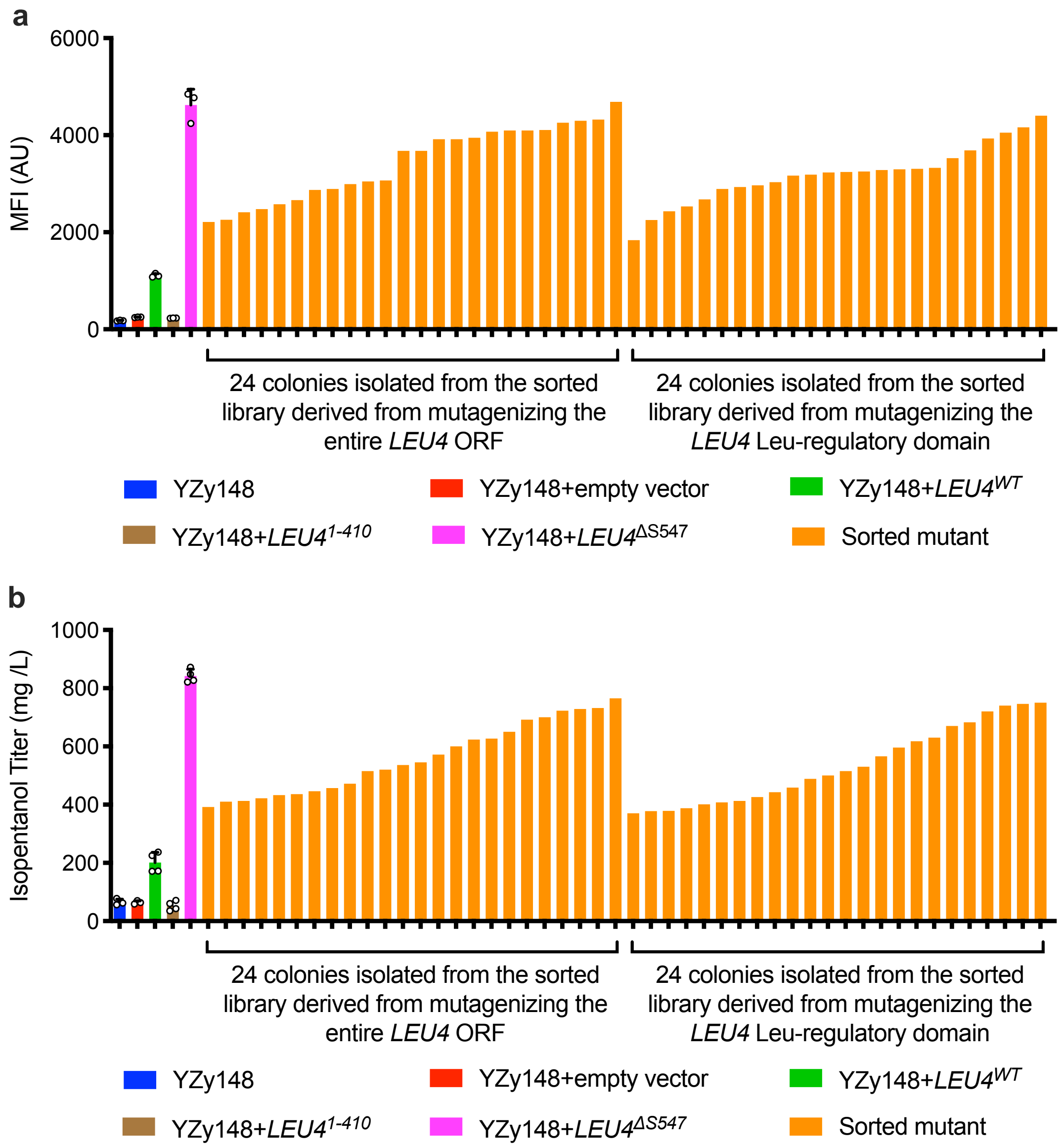

**Supplementary Figure 6. Screen of 48 random colonies sorted from libraries derived from mutagenizing the entire *LEU4* ORF or the *LEU4* regulatory domain.** (**a**) Flow cytometry measurements of the GFP median fluorescence intensity (MFI) of 48 sorted strains (orange) measured after 13 h of growth in SC-Ura media supplemented with 2% glucose (24 colonies are isolated from the sorted library derived from mutagenizing the entire *LEU4* ORF, and 24 colonies are isolated from the sorted library derived from mutagenizing the *LEU4* regulatory domain). MFI of the basal strain YZy148 (*leu4Δ leu9Δ bat1Δ LEU2*) with (red) or without (blue) an empty vector, or transformed with plasmids containing wild-type *LEU4* ($LEU4^{WT}$, green), $LEU4^{1\text{-}410}$ (brown), or $LEU4^{\Delta S547}$ (magenta), are shown as controls. (**b**) Isopentanol titers after 48h fermentations with the same 48 sorted strains (orange) and controls (red, blue, green, brown, and magenta) shown in (**a**). Error bars of the control strains represent the standard deviation of at least three biological replicates.

**Supplementary Figure 7**

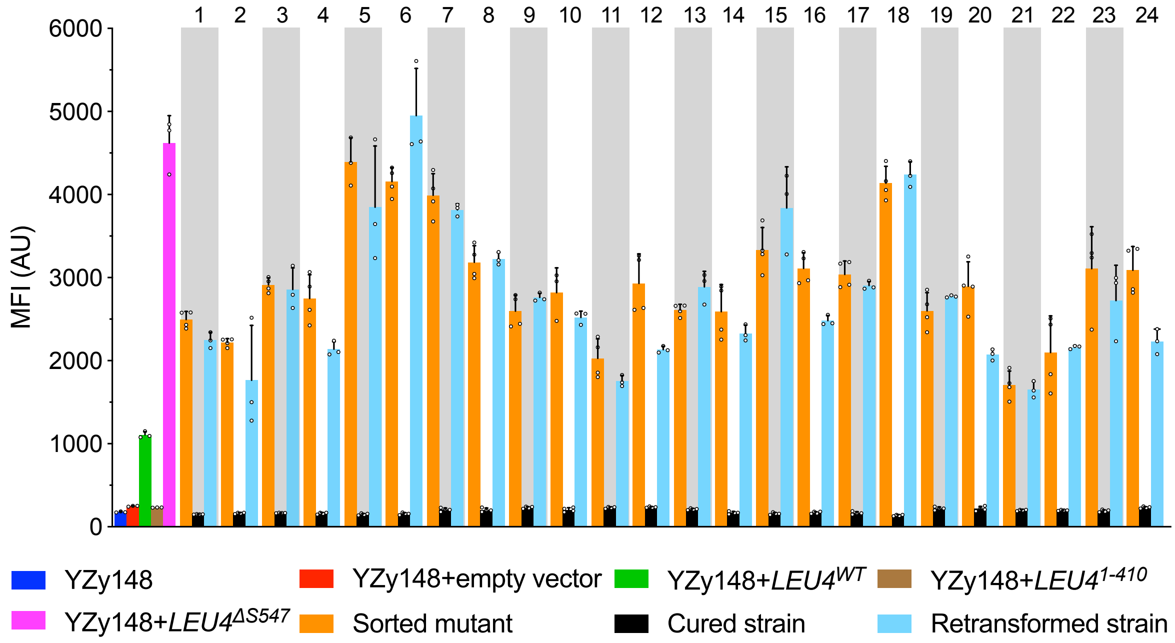

**Supplementary Figure 7. Confirmation that isolated *LEU4* variants with unique sequences enhance GFP fluorescence signal from the isopentanol biosensor.** Flow cytometry measurements of the GFP median fluorescence intensity (MFI) of sorted strains with unique *LEU4* sequences (orange), plasmid-cured derivatives (black), and strains obtained from retransforming YZy148 with each unique plasmid isolated from the corresponding sorted strains (cyan), measured after 13 h of growth in SC-Ura media supplemented with 2% glucose. GFP fluorescence intensity of the basal strain YZy148 (*leu4Δ leu9Δ bat1Δ LEU2*) with (red) or without (blue) an empty vector, or transformed with plasmids containing wild-type *LEU4* ($LEU4^{WT}$, green), $LEU4^{1\text{-}410}$ (brown), or $LEU4^{\Delta S547}$ (magenta), are shown as controls. The numbers on the upper x-axis identify each of the strains harboring screened mutants with unique *LEU4* sequences. *LEU4* mutants 1–13 were isolated from the sorted library derived from mutagenizing the entire *LEU4* ORF. *LEU4* mutants 14–24 were isolated from the sorted library derived from mutagenizing the *LEU4* leucine regulatory domain. Error bars represent the standard deviation of three biological replicates.

**Supplementary Figure 8**

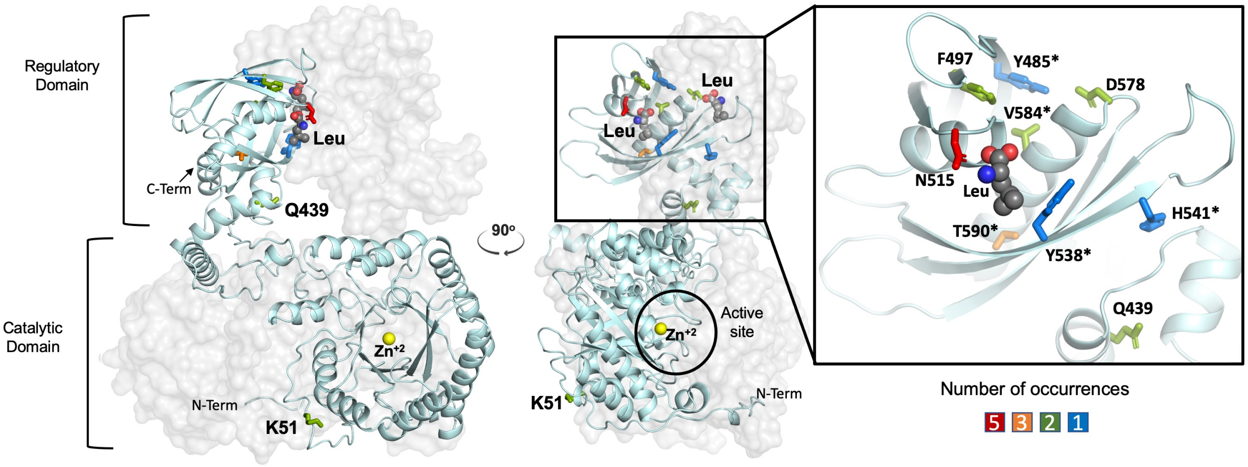

**Supplementary Figure 8. Structural analysis of key residues mutated in isolated Leu4p variants.** Key residues found to be mutated in Leu4p variants that enhance isopentanol production mapped onto the crystal structure of LeuA from *Mycobacterium tuberculosis* (pdb: 3fig). The catalytic and regulatory domains of the LeuA dimer are depicted (left panel). One of the two monomers is represented as a cartoon (light blue) and the other one as a surface (gray). A close-up view of the occupied leucine binding site of the regulatory domain (right panel box) shows the orientation of key residues (shown in sticks) found to be mutated in isolated *LEU4* variants that enhance isopentanol production. Residues labeled with an asterisk (*) are those mutated in variants containing only one mutation (see **Supplementary Table 5**). The number of unique sequences in which each residue is found mutated is depicted by a color scale shown in the bottom right corner (see **Supplementary Tables 4, 5**). Leucine bound to the regulatory binding site and a zinc ion are show in spheres.

**Supplementary Figure 9**

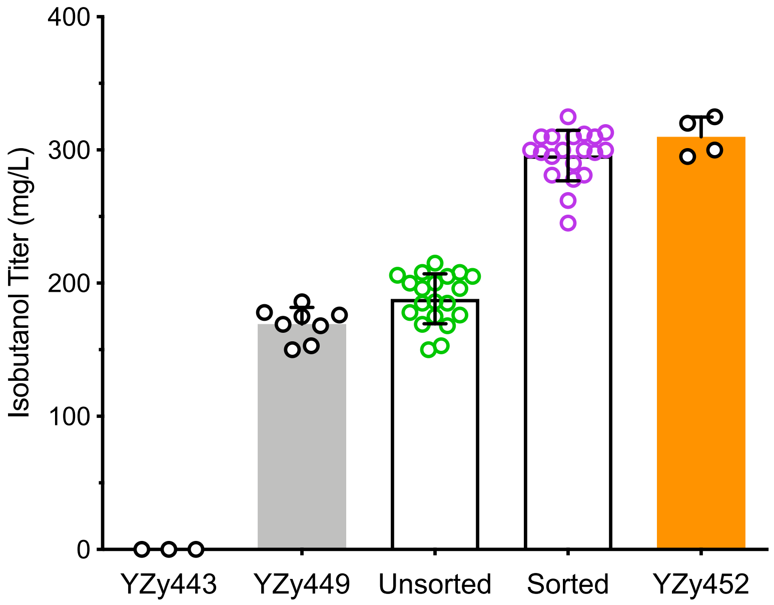

**Supplementary Figure 9. High-throughput screens, enabled by the isobutanol biosensor, to identify strains carrying extra copies of *Ec_ilvC*$^{P2D1\text{-}A1}$ with enhanced isobutanol production from galactose.** Isobutanol production of the parent strain (YZy443), the baseline strain containing a single copy of a genomically integrated galactose-inducible isobutanol cytosolic pathway (YZy449), 20 random colonies from each the unsorted population (unsorted) and sorted population (sorted) of YZy449 transformed with extra copies of *Ec_ilvC*$^{P2D1\text{-}A1}$ randomly integrated into δ-sites, and the best sorted strain (YZy452). Measurements were made after 48h fermentations in 15% galactose. Error bars represent the standard deviation of at least three biological replicates (for YZy443, YZy449 and YZy452) or 20 random colonies (from unsorted and sorted populations).

**Supplementary Figure 10**

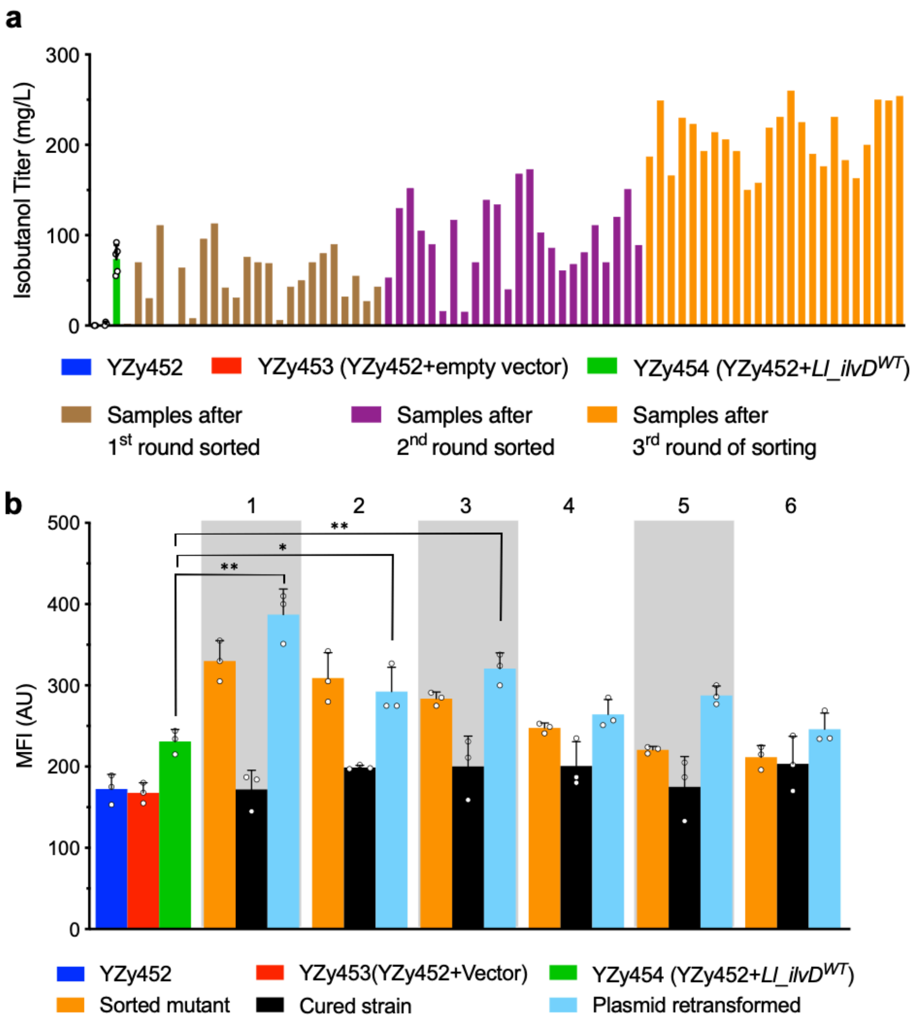

**Supplementary Figure 10. High-throughput screens for *Ll_ilvD* variants with enhanced cytosolic isobutanol production from glucose using the isobutanol biosensor.** (**a**) Isobutanol titers of 24 colonies randomly picked after each round of FACS. Titers obtained with the basal strain YZy452 (containing the cytosolic isobutanol pathway with extra copies of *Ec_ilvC*$^{P2D1-A1}$ and galactose-inducible *Ll_ilvD*) with (red) or without (blue) an empty vector, or transformed with a plasmid containing wild-type *Ll_ilvD* (*Ll_ilvD*$^{WT}$, green) are shown as controls. (**b**) Confirmation that isolated *Ll_ilvD* variants with unique sequences enhance GFP fluorescence signal from the isobutanol biosensor. Flow cytometry measurements of the GFP median fluorescence intensity (MFI), taken after 13h of growth in 2% glucose, for sorted strains (orange), plasmid-cured derivatives (black), and basal strain (YZy452) retransformed with each unique plasmid isolated from the corresponding sorted strains (cyan). Fluorescence of the basal strain YZy452 with (red) or without (blue) an empty vector, or transformed with a plasmid containing *Ll_ilvD*$^{WT}$ (green) are shown as controls. The numbers on the upper x-axis identify each of the strains harboring unique *Ll_ilvD* variants. Error bars represent the standard deviation of three biological replicates. A two-tailed *t*-test was used to determine the statistical significance of the difference between MFI of YZy452 transformed with a plasmid containing *Ll_ilvD*$^{WT}$ (green), or *Ll_ilvD* mutant #1

($Ll\_ilvD^{I433V}$) , or mutant #2 ($Ll\_ilvD^{V12A,\ S189P,\ H439R}$), or mutant #3 ($Ll\_ilvD^{K535R}$). * $P \leq 0.05$, ** $P \leq 0.01$.

**Supplementary Figure 11**

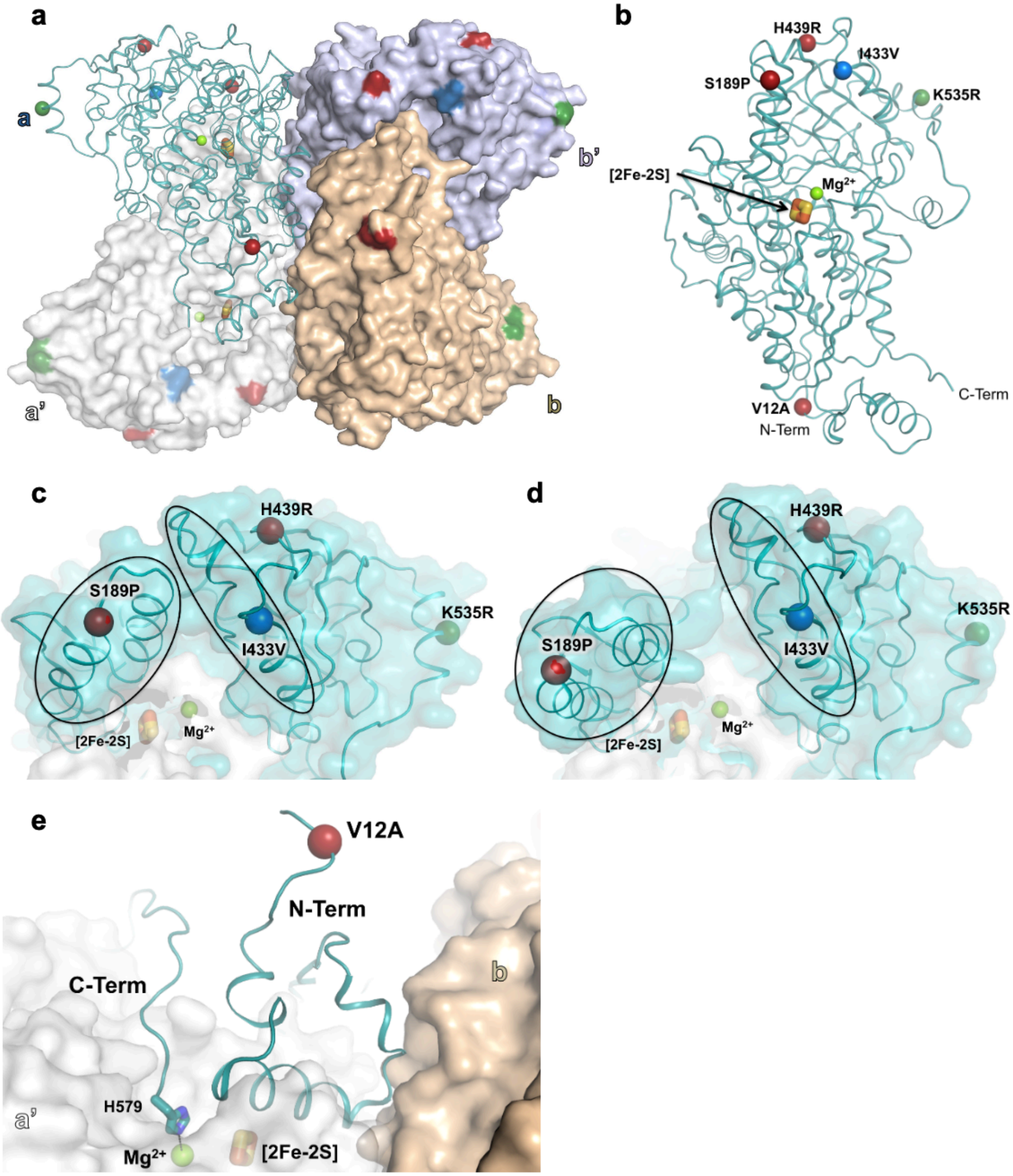

**Supplementary Figure 11. Structural analysis of key residues mutated in isolated IlvD variants.** Key residues found to be mutated in *Ll*_IlvD variants that enhance cytosolic isobutanol production mapped onto the crystal structure of its homolog l-arabinonate dehydratase from *Rhizobium leguminosarum bv. trifolii* (pdb: 5j84). The colored spheres indicate the locations of the mutated residues found in *Ll*_IlvD mutants #1 (blue), #2 (red) and #3 (green), (see **Supplementary Table 7**) (**a**) Tetramer of l-arabinonate dehydratase, a homolog of *Ll*_IlvD with 31.56% sequence identity and 36% sequence similarity. Residues substituted in *Ll*_IlvD mutants #1 (blue), #2 (red), and #3 (green) are shown as spheres on one of the four monomers represented as a ribbon (light blue), and colored on the surface representations (light blue, wheat, gray) of the other three monomers. (**b**) Ribbon representation of an *Ll*_IlvD monomer showing the positions of mutated residues (spheres) found in *Ll*_IlvD mutants #1 (blue), #2 (red) and #3 (green), relative to the 2Fe-2S cluster (orange and yellow sticks) and the $Mg^{2+}$ ion (bright green sphere) bound to the active site. (**c, d**) Close-up views of the closed (**c**) and open (**d**) conformations of the enzyme (pdb: 5j84 and pdb: 5j85, respectively). Residues S189 (substituted in mutant #2) and I433 (substituted in mutant #1), are located in the lobes (circled in black) that open and close to grant access or protect to the active site. (**e**) A close-up view shows the packing between the N- and C-termini of a monomer, which potentially contributes to the positioning of the His-579 that coordinates the $Mg^{2+}$ in the active site (found in a different monomer).

**Supplementary Figure 12**

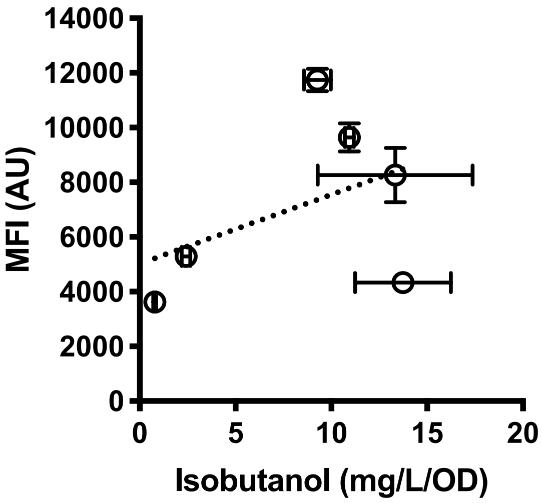

**Supplementary Figure 12. Correlation between specific isobutanol titers and GFP fluorescence signals from the isopentanol biosensor in the *LEU2* strains engineered for isopentanol production.** Strains used from left to right: SHy187, SHy192, SHy158, SHy176, SHy159, and SHy188 (**Supplementary Table 1**). The median fluorescence intensity (MFI) for each strain was measured after 13h of growth and plotted with the corresponding specific isobutanol titers obtained after 48h high-cell-density fermentations. The dotted line shows a poor linear regression fit with an $R^2$ of 0.19. Error bars represent the standard deviation of at least three biological replicates.

**Supplementary Figure 13**

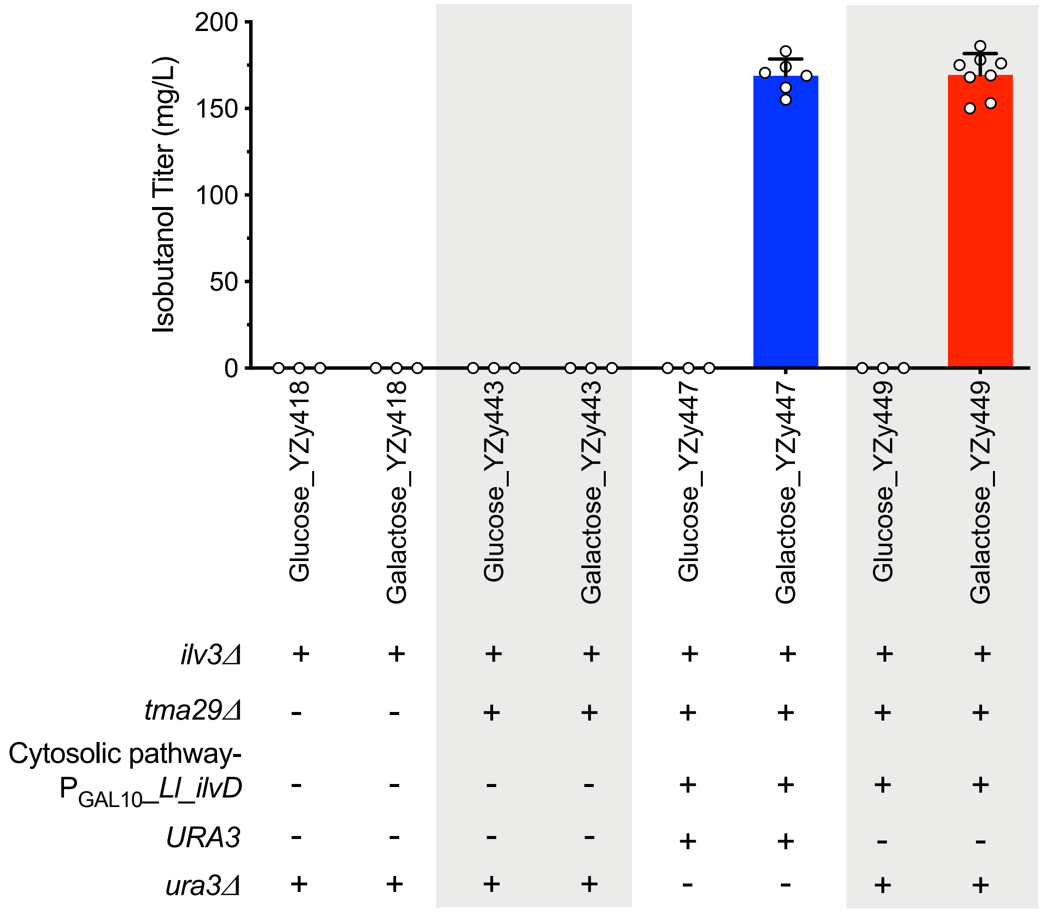

**Supplementary Figure 13. Isobutanol production from glucose and galactose in engineered strains with or without galactose-inducible *Ll_ilvD*.** Strains were fermented for 48h using 15% glucose or galactose as the carbon source. YZy418 and YZy443 produced no detectable isobutanol from either carbon source. Error bars represent the standard deviation of at least three biological replicates.

1. Entian, K.-D. & Kötter, P. 25 Yeast Genetic Strain and Plasmid Collections. *Methods in Microbiology* **36**, 629-666 (2007).
2. Hammer, S.K. & Avalos, J.L. Uncovering the role of branched-chain amino acid transaminases in Saccharomyces cerevisiae isobutanol biosynthesis. *Metabolic Engineering* **44**, 302-312 (2017).
3. Sikorski, R.S. & Hieter, P. A system of shuttle vectors and yeast host strains designed for efficient manipulation of DNA in Saccharomyces cerevisiae. *Genetics* **122**, 19 (1989).
4. Christianson, T.W., Sikorski, R.S., Dante, M., Shero, J.H. & Hieter, P. Multifunctional yeast high-copy-number shuttle vectors. *Gene* **110**, 119-22 (1992).
5. Zhao, E.M. et al. Optogenetic regulation of engineered cellular metabolism for microbial chemical production. *Nature* **555**, 683-687 (2018).
6. Zhang, Y. et al. Xylose utilization stimulates mitochondrial production of isobutanol and 2-methyl-1-butanol in Saccharomyces cerevisiae. *Biotechnol Biofuels* **12**, 223 (2019).
7. Zhao, E.M. et al. Design, characterization, and modeling of rapid optogenetic inverter circuits for dynamic control in yeast metabolic engineering, Under review. (2020).
8. Goldstein, A.L. & McCusker, J.H. Three new dominant drug resistance cassettes for gene disruption in Saccharomyces cerevisiae. *Yeast* **15**, 1541-53 (1999).
9. Avalos, J.L., Fink, G.R. & Stephanopoulos, G. Compartmentalization of metabolic pathways in yeast mitochondria improves the production of branched-chain alcohols. *Nat Biotechnol* **31**, 335-41 (2013).
10. Gueldener, U., Heinisch, J., Koehler, G.J., Voss, D. & Hegemann, J.H. A second set of loxP marker cassettes for Cre-mediated multiple gene knockouts in budding yeast. *Nucleic Acids Res* **30**, e23 (2002).
11. Atsumi, S., Li, Z. & Liao, J.C. Acetolactate synthase from Bacillus subtilis serves as a 2-ketoisovalerate decarboxylase for isobutanol biosynthesis in Escherichia coli. *Appl Environ Microbiol* **75**, 6306-11 (2009).
12. Brinkmann-Chen, S. et al. General approach to reversing ketol-acid reductoisomerase cofactor dependence from NADPH to NADH. *Proc Natl Acad Sci U S A* **110**, 10946-51 (2013).
13. Urano, J.E., CO, US), Dundon, Catherine Asleson (Englewood, CO, US). Cytosolic isobutanol pathway localization for the production of isobutanol. (Gevo, Inc. (Englewood, CO, US), United States, 2012).
14. Kohlhaw, G.B. Leucine Biosynthesis in Fungi: Entering Metabolism through the Back Door. *Microbiology and Molecular Biology Reviews* **67**, 1-15 (2003).
15. Hazelwood, L.A., Daran, J.M., van Maris, A.J.A., Pronk, J.T. & Dickinson, J.R. The ehrlich pathway for fusel alcohol production: a century of research on Saccharomyces cerevisiae metabolism. *Applied and Environmental Microbiology* **74**, 2259-2266 (2008).
